# Phylogenomic synteny analysis tracks conserved ancient polyploid-derived triplicated genomic blocks across Asteraceae genomes

**DOI:** 10.1101/2025.01.08.631874

**Authors:** Tao Feng, Michael McKibben, John Lovell, Richard Michelmore, Loren H. Rieseberg, Michael Barker, Eric M. Schranz

**Author notes:** To whom correspondence should be addressed: Eric Schranz. **Email:**.

## Abstract

The Asteraceae (Compositae) is the largest flowering plant family, ubiquitous in most terrestrial communities, and morphologically hyper-diverse. An ancient whole genome triplication (paleo-hexaploidization) occurred at approximately the same time as the evolutionary innovation and adaptive radiation of the family during the middle Eocene. Despite its importance, the genomic contents arising from this triplication have yet to be tracked in context of the Asteraceae genome evolution. We applied a synteny oriented phylogenomic analysis of 21 Asterales genomes and to study the paleo-hexaploidization and its consequences to gene, trait, and genome evolution. We identified 15 ancestral linkage groups (ALGs) that date back to the common diploid ancestor of all Asteraceae. Each of these groups was triplicated, resulting in 45 genomic blocks (3×15), which serve as the foundation for cross-family analyses. We demonstrate the complex evolutionary dynamics of the 45 genomic blocks across the Asteraceae phylogeny. We found that modern genomes are genetic mosaics of three progenitor genomes by extensive genomic exchange, chromosomal shuffling and gene fractionation. 157 genes retained three paleo-hexaploid derived syntenic paralogs across most Asteraceae species. Transcription factors (TFs) and auxin-related genes are significantly overrepresented in the conserved triplets, and expression of the paleo-hexaploidy paralogs is spatiotemporally differentiated. These genes are involved in the development of floral capitulum, a remarkable morphological innovation of the family. The discovery of conserved triplicated genes can direct further study to understand the evolutionary innovation, and the synteny-phylogenomic framework and ALGs provide a comparative framework to characterize newly sequenced Asteraceae genomes.

## Introduction

The Asteraceae, traditionally known as Compositae, is composed of more than 34,000 species, corresponding to ∼10% of all flowering plants (https://wfoplantlist.org/; accessed at 12/12/2024). A shared derived feature of all Asteraceae species is the capitulum, a head-like inflorescence mimicking a single flower, which is perhaps the most remarkable morphological innovation in angiosperms after the origin of flowers (1). The Asteraceae are exceptionally diverse in phenotypic traits, specialized metabolites, and ecological habitats, making this family an excellent system to address a broad range of eco-evolutionary questions (2). Asteraceae have colonised almost every conceivable habitat including the harshest ones, for example *Saussurea gnaphalodes* is endemic to up to 6400m in the Himalayan periglacial region, the uppermost elevation limit recorded for vascular plants (3). There are also species such as the common sunflower (*Helianthus annuus*) and dandelion (*Taraxacum officinale*) that are widely distributed across the globe. The family is well known for its agriculturally important species such as cultivated sunflower, safflower, lettuce, and globe artichoke and iconic horticultural species such as daisies, gerbera and chrysanthemums. In addition to their vast morphological and habitat diversity, Asteraceae harbour extensive genomic variation with ploidy levels ranging from 1x to 22x (2n=4 to198) and 2C-values varying 139-fold, making it more diverse than most other angiosperm families (4). The hyper-diversity of Asteraceae is suggested to be generated by multiple radiation events during the middle Eocene (5).

Despite the high diversity and species richness of Asteraceae, genome sequencing the diversity of the family has lagged, with most genome sequences being generated in the past three years. Sequencing and assembly of Asteraceae genomes have been challenging because these genomes are generally large and consist of long and highly similar repeats (6). However, technical advances in long-read sequencing and bioinformatics have overcome many of these obstacles, bringing about a new era of comparative genomics for the family. The first complete genome sequence of the Asteraceae was for the globe artichoke (*Cynara cardunculus* var. *scolymus*) (7), followed by the sequencing of sunflower (*Helianthus annuus*) (6) and lettuce (*Lactuca sativa*) (8). As of the time of our analysis, a total of 61 assemblies from 35 distinct species were publicly available (https://www.ncbi.nlm.nih.gov/datasets/genomes/?taxon=4210). There will certainly be an exponential growth of genome sequencing of Asteraceae species in the coming years, highlighting the need for a unified and consistent comparative genomics framework for the family.

Taxonomic and phylogenetic studies of Asteraceae have been challenging, both due to the extremely high species richness but also due to widely occurring hybridization, polyploidization and rapid radiations (9, 10). However, recent advances in phylogenomic approaches and high-throughput sequencing (including genome, transcriptome and targeted capture sequencing) have enabled significant advances in clarifying phylogenetic relationships in Asteraceae (5, 10–13). Recent phylogenomic studies (5, 13) have reached a near-consensus phylogenetic backbone for the family (Fig. S1), with only minor inconsistencies in the positions of several small tribes, likely because of putative gene tree conflicts. Aligning with this backbone, the phylogenetic classification of Asteraceae was recently updated (14), which includes 16 subfamilies and 47 tribes (or 51 tribes if several evolutionary clades are treated as individual tribes). A recent phylogenomic study found Barnadesioideae is closer to Calyceraceae (15), making Asteraceae a potentially polyphyletic group.

In addition, phylogenomic studies and genome sequencing have demonstrated widely occurring whole genome duplications (WGD) across the Asteraceae phylogeny, including ancient events during early Asteraceae evolution (12, 13, 16, 17) and more recent events in specific lineages (6, 12, 13, 18, 19). According to these studies, Asteraceae has undergone two successive ancient polyploidization events, the first one shared by Asteraceae and Calyceraceae and the second one shared by all Asteraceae species except for the two first-diverging subfamilies (17, 20). In genomic content, Calyceraceae, Barnadesioideae and Famatinanthoideae are paleo-tetraploids, while all other extant Asteraceae are paleo-hexaploids. This two-step hexaploidization was suggested to be a key event in Asteraceae evolution, which might be the main driver of the rise of Asteraceae during the middle Eocene (16, 17). However, the contribution of this hexaploidization to the architecture and content of modern Asteraceae genomes is poorly understood. Nonetheless, a robust phylogenetic backbone, and the scenarios of genome duplication in Asteraceae provide a useful framework for comparative genomic study.

In this study, we aimed to illustrate and understand Asteraceae genome evolution in the context of the paleo-hexaploidization. We established the first synteny constrained phylogenomic framework for the family using 20 high-quality genomes, from diploid (2n=2x) to paleotraploid (2n=24x) species and included *Scaevola taccada* (Goodeniaceae) as the non-triplicated outgroup. Using this framework, we aimed to answer: 1) to what extent can we trace the genome changes across ∼80 million years of Asteraceae evolution?; 2) what is the genomic architecture of modern Asteraceae genomes compared with the inferred non-triplicated ancestral genome?; 3) what genomic rearrangements occurred during the evolution of the extant paleo-polyploid genomes relative to the inferred progenitor genome?; 4) what is the consequence of genome multiplication on phenotypic trait evolution? Answers to these questions provided insights into Asteraceae genome evolution and its rise to ecological prevalence.

## Results

### Macro- and micro-synteny across Asteraceae genomes

To build a synteny phylogenomic framework for Asteraceae, we gathered 30 chromosome-level genome assemblies of Asteraceae (Fig. S1, Table S1). In addition, we also included one outgroup genome from the sister family Goodeninaceae. We assessed the quality of the genome data using the widely used BUSCO (21) and a recently developed phylogeny-aware approach OMArk (22) which use a predefined set of conserved orthologous groups as a proxy for genome-wide completeness. OMArk gave slightly different but consistent assessments of the genome assemblies (Table S1). According to the assessment, taking phylogenetic representation into account, 20 genome assemblies (OMArk: 90.36-98.12; BUSCO: 90.4-98.2) representing 10 tribes in three Asteraceae subfamilies and the outgroup Goodeniaceae were used for comparative analyses.

With this dataset, we built the first genome synteny map across the Asteraceae and its sister family Goodeninaceae, spanning >80 million years of evolutionary divergence (Fig. 1a). Despite the divergence, there was extensive genome synteny across species (Fig. 1a). The overall synteny map captured the ancient polyploid events, as seen by the ratios of syntenic regions across the phylogeny. This includes the ancient polyploidization shared by all Asteraceae species, the ancient duplication shared by *H. annuus*, *Scalesia atractyloides*, *Mikania micrantha* and *Smallanthus sonchifolius*, and the recent lineage-specific duplications in *S. atractyloides* and *S. sonchifolius* (Fig. 1a). Our results support the previous observations based on sequence divergence (KS frequency) and gene tree sorting (12, 17).

**Fig. 1.**
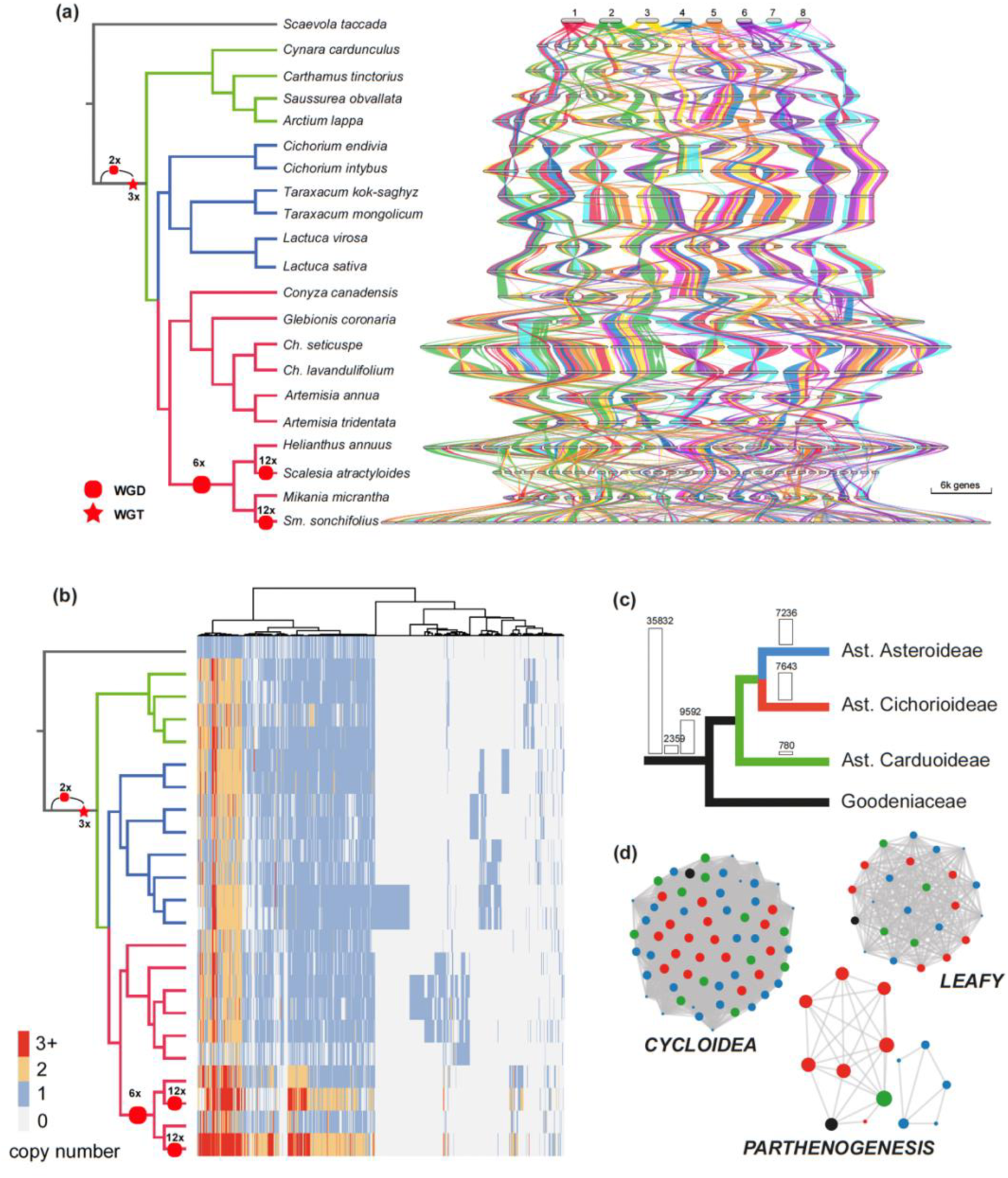
Macro- and Micro-synteny across Asteraceae and Goodeninaceae genomes. **a.** Macrosynteny synteny map (riparian plot) of orthologous regions among 21 genomes, with the phylogeny adapted from Zhang et al. (13) and the scenario of WGD labelled in nodes. Chromosomes are ordered horizontally to maximize synteny with *Scaevola taccada* and ribbons are colour coded by synteny to *S. taccada* chromosomes; **b.** Phylogenomic synteny profiling of micro-synteny gene clusters (size ≥ 2) across 21 genomes. Rows are species in same order as in a, and columns are synteny clusters each of which is an assemblage of homologous genes that are syntenic across two or more species. Gene copy number is indicated by colour bars. The presence/absence pattern is clustered using Euclidean distance, as indicated by the tree on top; **c.** Quantitative characteristics of micro-synteny clusters. On the root stem is the total number of synteny cluster, conserved synteny clusters across Asteraceae and Goodeninaceae, and conserved synteny clusters within Asteraceae, respective. On the tip branches are the lineage-specific clusters in three subfamilies; **d.** Network visualization of different type of synteny clusters, *CYCLOIDEA* (multiple-copy), *LEAFY* (single-copy) and *PARTHENOGENESIS* (sub-clusters by translocation). Nodes are gene with colour indicating their phylogenetic affiliation showed in c, and edges are syntenic relations between genes.

Based on this macro-synteny framework, we then examined the micro-synteny namely the synteny of individual genes across Asteraceae. We identified 35,832 synteny clusters (a group of orthologs that are syntenic across two or more genomes (23)), of which 2,359 and 9,592 are highly conserved across Asteraceae and Goodeninaceae, and within Asteraceae respectively (Fig. 1b-c). These clusters comprised related genes that are present and syntenic across the phylogeny after over 80 million years of genome shuffling, gene fractionation, and transposition. Therefore, they are the core syntenic genes of the Asteraceae. We also detected 7,236, 7,643, and 780 lineage-specific synteny clusters in the subfamilies Asteroideae, Cichorioideae, and Carduoideae, respectively (Fig. 1c).

Because of genome duplication, gene fraction, and translocation, a synteny cluster could be a multi-copy or single-copy cluster, and/or may be divided into multiple sub-clusters, as exemplified by gene *CYCLOIDEA*, *LEAFY* and *PARTHENOGENESIS* respectively (Fig. 1d). Our result also highlights the Asteroideae species with more gene copies per cluster (e.g., orange/red rows in Fig. 1b), which is consistent with the observed additional WGD in these species (Fig. 1a).

### Genomic architecture of Asteraceae genomes in context of paleo-hexaploidization

One of our main objectives is to understand the consequence of the ancient two-step hexaploidization (ambiguously termed as triplication elsewhere, e.g. (6, 8, 24)) on genome evolution in Asteraceae. To characterize the genome architecture of extant Asteraceae species in context of the paleo-hexaploidization, we catalogued Asteraceae genomes into genomic blocks. Using *Scaevola taccada* as the outgroup, we focused on diploid Asteraceae species and quantified the genome shuffling events from *S. taccada* to Asteraceae species. *Scaevola taccada* was used as an outgroup because it is a close sister to Asteraceae but does not share the paleo-hexaploidization and has no duplication after its divergence with Asteraceae (24). On average, 152 fissions and 165 fusions are identified in pairwise comparisons (Fig. S2). The *Arctium lappa* genome is the least rearranged among the genomes analyzed, with 75 inferred fissions and 81 inferred fusions (Fig. S2). In addition, the *A. lappa* genome assembly is of high quality in both completeness (BUSCO=98, OMArk=97.76) and continuity (LAI=21.57, gold quality according to the classification based on LTR Assembly Index (25)). Therefore, *A. lappa* was used as an in-group reference to compare with the outgroup *S. taccada* to define Asteraceae genomic blocks.

Using the A-Bruijn graph-based algorithm (26), we screened conserved syntenic segments between *A. lappa* and *S. taccada*. In total, 1324 conserved syntenic segments (at least 6 genes) were identified in *A. lappa*, of which 291, 460, and 573 segments are in triple, double, and single states (Fig. S3), and the largest one has 97 genes (15 genes on average). Mapping the 1,324 syntenic segments onto the 18 *A. lappa* chromosomes indicated that the *A. lappa* genome is a mosaic of three progenitor genomes (Fig. 2a), as exemplified by chromosome 14 and 17 (homoeologous to chromosome 5 of *S. taccada*), and their paralogous segments are widely distributed across chromosome 1 (Fig. 2a, Fig. S3). Therefore, most extant Asteraceae genomes, like *A. lappa*, are mosaics of the three progenitor genomes.

**Fig. 2.**
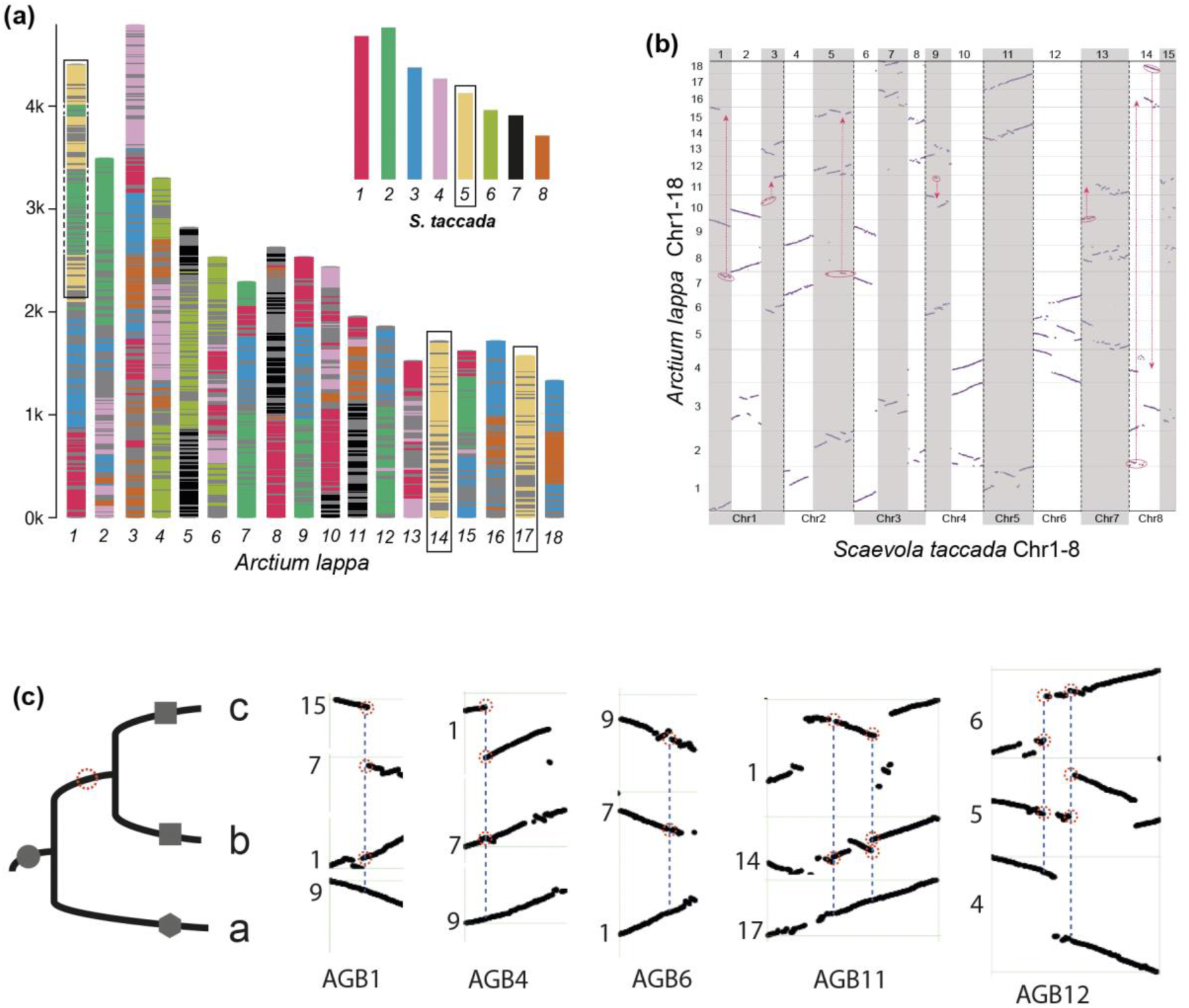
Genomic comparison between *Scaevola taccada* and *Arctium lappa* and the 15 Asteraceae genomic blocks (AGBs). **a.** The mosaic architecture of *A. lappa* genome with genomic segments coloured to show homology to *S. taccada* chromosomes. As an example, *S. taccada* chromosome 5 is homology to *A. lappa* chromosome 14, 17 and top parts of chromosome 1. The chromosome length indicates number of genes rather than nucleotide base pairs; **b.** Dot plot between *S. taccada* (x-axis) and *A. lappa* (y-axis) genome. The eight chromosomes of *S. taccada* are divided into 15 regions each aligning with three regions on *A. lappa* chromosomes. The inferred genomic fission/fusion events across different chromosomes of *A. lappa* are highlighted in circles and arrows; **c.** For AGB1, 4, 6, 11, 12 the subgenome a can be inferred according to the principle of parsimony, i.e., the breakage shared by two homoeologous blocks (in red dotted circle) is likely a single event occurred in their common ancestral block rather than two events happened independently.

Pairwise syntenic comparisons between *S. taccada* and *A. lappa* showed a clear syntenic depth ratio of 1:3 (Fig. 2b), and considerable part of the *A. lappa* genome including the whole chromosome 14 and 17, have retained to be intact with only several inversions compared to *S. taccada* genome (Fig. 2b). This enables us to identify the homoeologous genomic regions that are presumably derived from the ancient genome polyploidization. Following the principles of proximity and complementarity, we assigned the identified genomic blocks of *A. lappa* to 15 sets of homoeologous blocks (blocks with common shared ancestry, Fig. 2b, Fig. S4), each of which contains three blocks that are inferred to be the descendants of the paleohexaploid ancestor (termed as AGB1-15 hereafter). For example, the chromosome 14, 17 and several segments on chromosome 1 constitute the AGB 11 which is homoeologous to the chromosome 5 of *S. taccada* (Fig. 2b-c). The homoeologous blocks in AGB14 cannot be fully assigned here but were resolved by cross-referring to the comparison between *S. taccada* and *Conyza canadensis* (Fig. S5). In the case of AGB 1, 4, 6, 11 and 12, the homoeologous blocks were phased to sub-genomes (Fig. 2c) based on the assumption that a rearrangement/breakage shared by two homoeologous blocks is likely a single event that occurred in their common ancestral block rather than two events that happened independently.

### Characterization of the divergence of subgenomes in Asteraceae

Next, we used the homoeologous blocks as units to explore the divergence of the subgenomes. To investigate whether there is gene fractionation bias among the three subgenomes, we calculated the gene retention rate in the homoeologous blocks in each AGB using *S. taccada* as a reference. Overall, there is no evidence of significant gene fractionation bias among subgenomes (Fig. 3a). As exemplified by AGB11 (Fig. 3b), the three descendants have ∼25%-55% gene retention rate, and no subgenome is clearly and consistently less fractionated, although some regions, for example the window300-400 in AGB11a, has more genes retained (Fig. 3b). In addition, we calculated the synonymous substitution rates (*Ks*) using syntenic genes and compared the *Ks* distributions among the homoeologous blocks. There is also no significant divergence in *Ks* distributions among the homoeologous blocks (Fig. S6), as exemplified by ABG11 (Fig. 3c-d). Further phylogenetic analysis using a sliding-window approach revealed extensive conflicts among the homoeologous AGBs and the outgroup *S. taccada* (Fig. 3e-f). For example, on AGB11, 34.7% windows support the topology *(o (a(b,c)))*, while 18.4% and 46.9% windows support alternative topologies *(o(b(a,c)))* and *(o (c(b,a)))*, respectively (Fig. 3d). These observations suggested that the three sister ABGs have undergone extensive genomic exchange in the paleohexaploid ancestor genome, and the exchange was shared by descendants of the paleohexaploid, including all species used in this study. The homogeneousness of the homoeologous genomic blocks hindered further subgenome phasing. Instead, we assigned the AGBs to three groups (a, b, c) randomly except for AGB 1, 4, 6, 11 and 12 for which subgenome a can be identified (Fig. 2c).

**Fig. 3.**
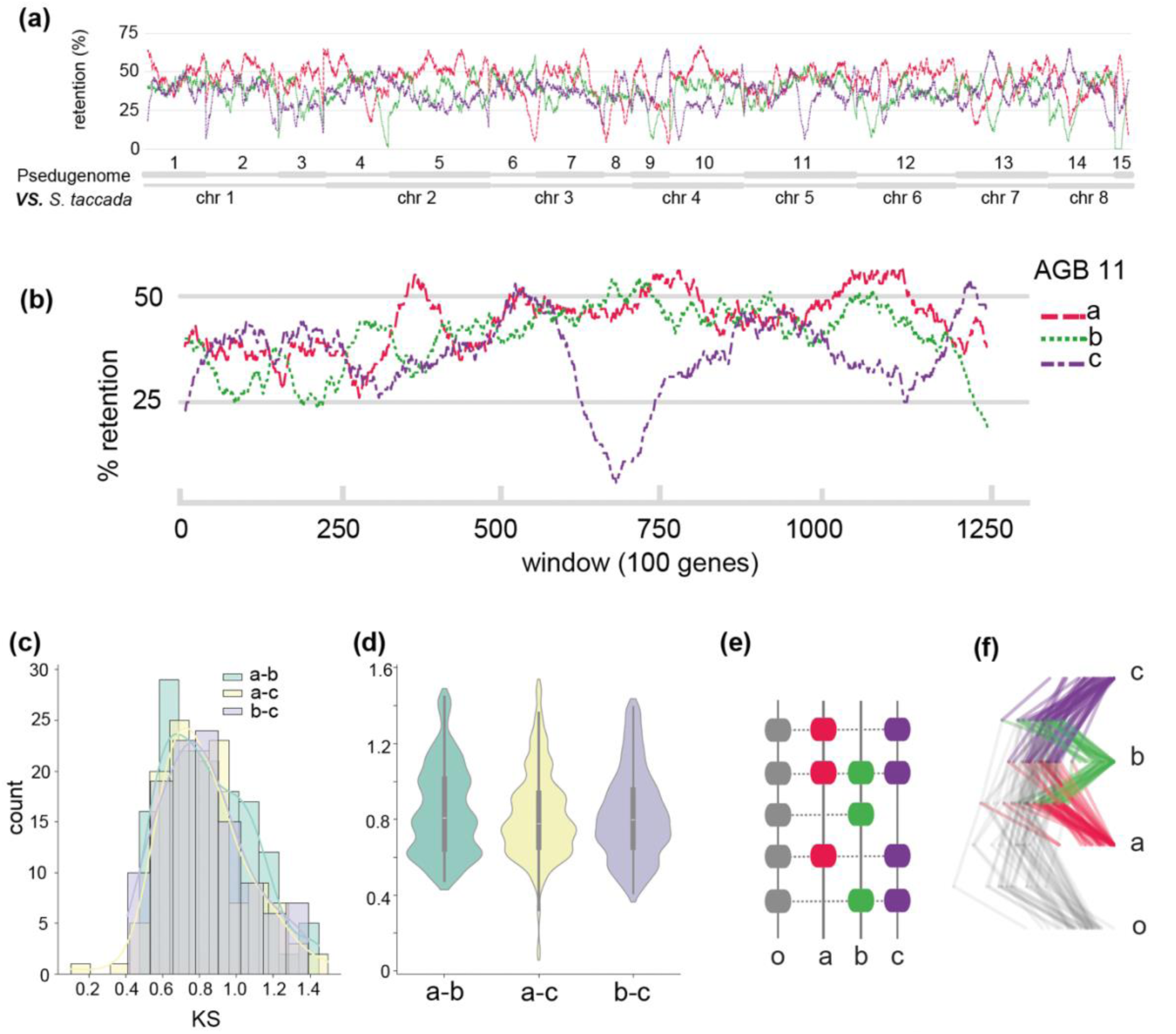
Characterization of the three subgenomes. **a.** The retention rate of syntenic genes in the three subgenomes; **b.** A zoom-in window of gene fractionation pattern of AGB11; **c-d.** Distribution of synonymous substitution rates (*Ks*) of syntenic genes in AGB11. The *Ks* plots for other AGBs are in Fig. S6; **e.** Diagram shows a window of syntenic genes from the subgenome a, b, c and outgroup o; **f.** Densitree of the phylogenies inferred from the syntenic genes in ABG11.

### A subgenome phased synteny-phylogenomic framework for Asteraceae

The 45 AGBs represent the likely genomic architecture and content of the paleohexaploid that is the common ancestor to most Asteraceae species. Based on *A. lappa* genome sequence, we generated a pseudo-genome with 15x3=45 pseudo-molecules (Fig. 4, Fig. S7). The pseudo genome is 1.52Gbp in length with 14,647, 12,205 and 13,135 genes on the three sets of ABGs (Fig. S7). It is important to clarify that the pseudo genome does not represent the Asteraceae ancestral genome directly, as *S. taccada* itself has likely undergone lineage-specific rearrangements since diverging from Asteraceae. However, given that *S. taccada* has not experienced any genome duplication events post-divergence and likely underwent fewer genome rearrangements compared to extant Asteraceae genomes, we propose that the pseudo genome provides a reasonable representation of the Asteraceae ancestral genome. Thus, we defined these reconstructed blocks as the ancestral linkage groups (ALGs) of Asteraceae.

**Fig. 4.**
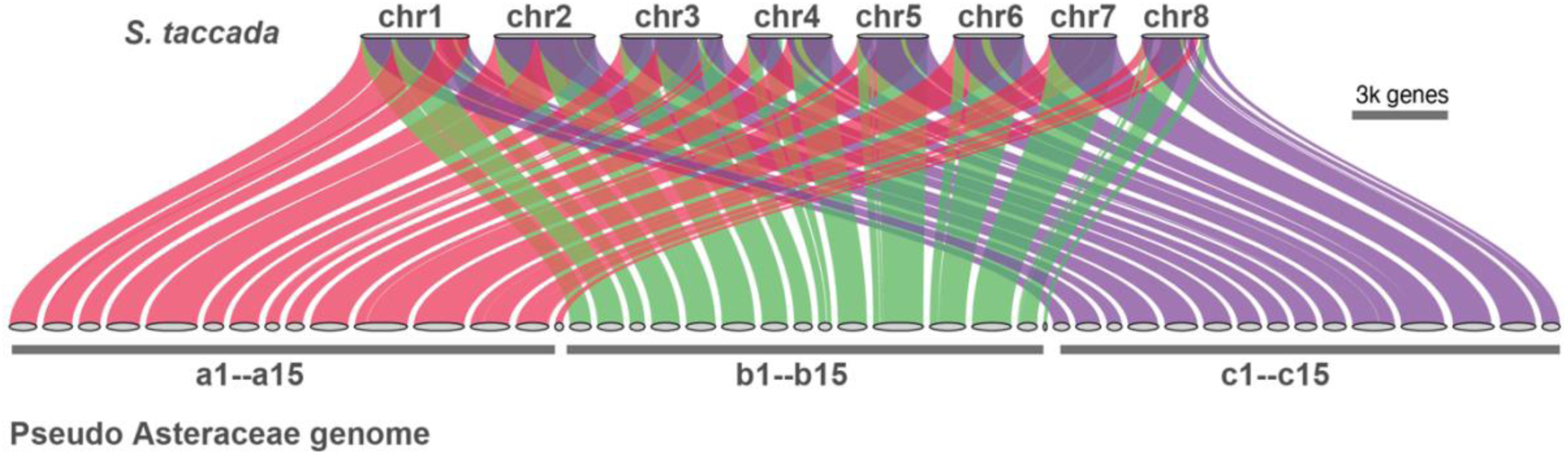
The sub-genome phased pseudo-genome. Macro-synteny between *S. taccada* genome and the pseudo genome with 15x3 pseudo chromosomes. The a,b,c represent three sets of subgenome derived from the two-step genome polyploidization in Asteraceae.

Next, we catalogued the Asteraceae genomes in the context of the 3x15=45 AGBs. We used the pseudo-genome as a reference and mapped the Asteraceae genomes onto it using the muti-genome synteny inference approach GENESPACE (27). With this, we generated an updated macrosynteny map across Asteraceae with the ancestral triplicated genomic blocks phased (Fig. 5, Fig. S8). Given any genomic region or genes of interest, for example the AGB11 (highlighted in Fig. 5a), the evolutionary dynamics can be traced across Asteraceae phylogeny. This catalogued genomic resource of 20 Asteraceae species and the outgroup *S. taccada*, including macrosynteny (Fig. S8) and microsynteny (Fig. S9), provides a framework to conduct comparative phylogenomic study in Asteraceae. We have developed a pipeline (https://github.com/xiaoyezao/Asteraceae-synteny-phylogenomics) for mapping genomes onto the 15x3 AGBs, therefore, newly generated genome sequences can be readily incorporated into this framework. To demonstrate the usage of this pipeline, we analysed the newly generated milk thistle genome (Fig. S10) (28) which was not used in our original analysis.

**Fig. 5.**
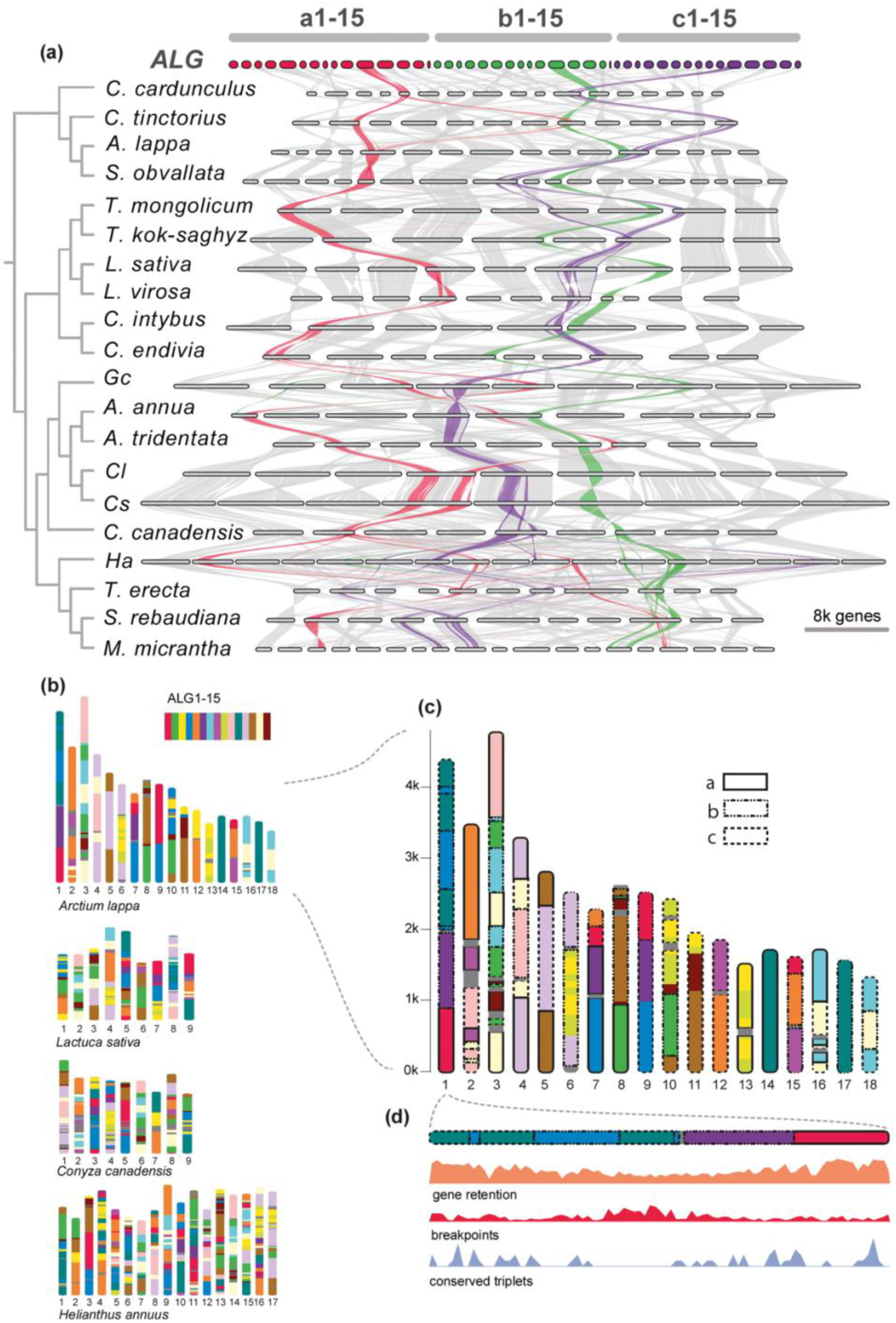
Sub-genome phased synteny-phylogenomic framework for Asteraceae. **a.** Macrosynteny synteny (riparian plot) across 20 Asteraceae genomes and the pseudo genome, with AGB11-a,b,c highlighted in red, green and purple colour to show the dynamics of AGB evolution in Asteraceae; **b.** The genome architecture of four representative species with genomic segments in different colours for tracking the inheritance of ALGs; **c.** Zooming in on the *Arctium lappa* genome to show the three paralogous regions derived from same AGB, which labelled in three types of rectangles; **d.** Zooming in on the chromosome 1 of *A. lappa* with the different genomic features (gene retention, chromosome breakpoints, and CTGs) represented in density along chromosomes by 100-gene window.

### 157 genes retain three syntenic paleo-hexaploid derived paralogs after fractionation

To further understand the consequence of polyploidization on gene content, we analysed gene fractionation, a process by which gene copies are eliminated from one homoeologous chromosome or the other(s) after polyploidization. Taking ABG11 as an example, 40%-80% genes (per 100-gene window) have been lost (Fig. 3b), and this is consistent across species (Fig. S11). In addition, gene fractionation along chromosomes behaves in a complementary pattern among the three homoeologous AGBs, i.*e.*, for a given region, high retention in one AGB supplement high fractionation in its homoeologous AGBs (Fig. S11), which results in, on average, a 124% gene retention rate (out of a possible 300% in total considering paleo-hexaploidy paralogs) in extant Asteraceae genomes.

The retention of 124% paleo-hexaploidy homologs in modern Asteraceae species indicates that gene fractionation has eliminated most paleo-paralogous copies. However, 24% paleo-paralogs have been retained after 80 million years of Asteraceae evolution. To independently validate patterns of fractionation we ran the machine learning tool Frackify (29) on 14 diploidy Asteraceae genomes. These analyses recovered that on average 23% of each genome were paleo-paralogous genes, and each species contained on average 860 triple-copy paleo-paralogs, consistent with our synteny based analyses (Table S2).

The triple-copy paleo paralogs identified in Asteraceae species are important genetic heritage of the ancient polyploidization, may contributed substantially to Asteraceae evolution. To further investigate the paleo paralogs in context of Asteraceae phylogeny, we performed multi-species comparison and found that 157 genes (471 paleo paralogs) were fully retained across all three Asteraceae subfamilies studied here (Fig. 6a, Table S3). It is likely that these genes are conserved across most Asteraceae lineages in terms of the retention of paleo paralogs because the sampling here covers diverse evolutionary lineages except for the basal ones (Fig. S1). These genes are termed as conserved triplicated genes (CTGs), hereafter. Further investigation of the CTGs indicates that they are distributed across the 15x3 AGBs (Fig. 6a) and form several hotspots, such as on AGB10 (Fig. 6a).

**Fig. 6.**
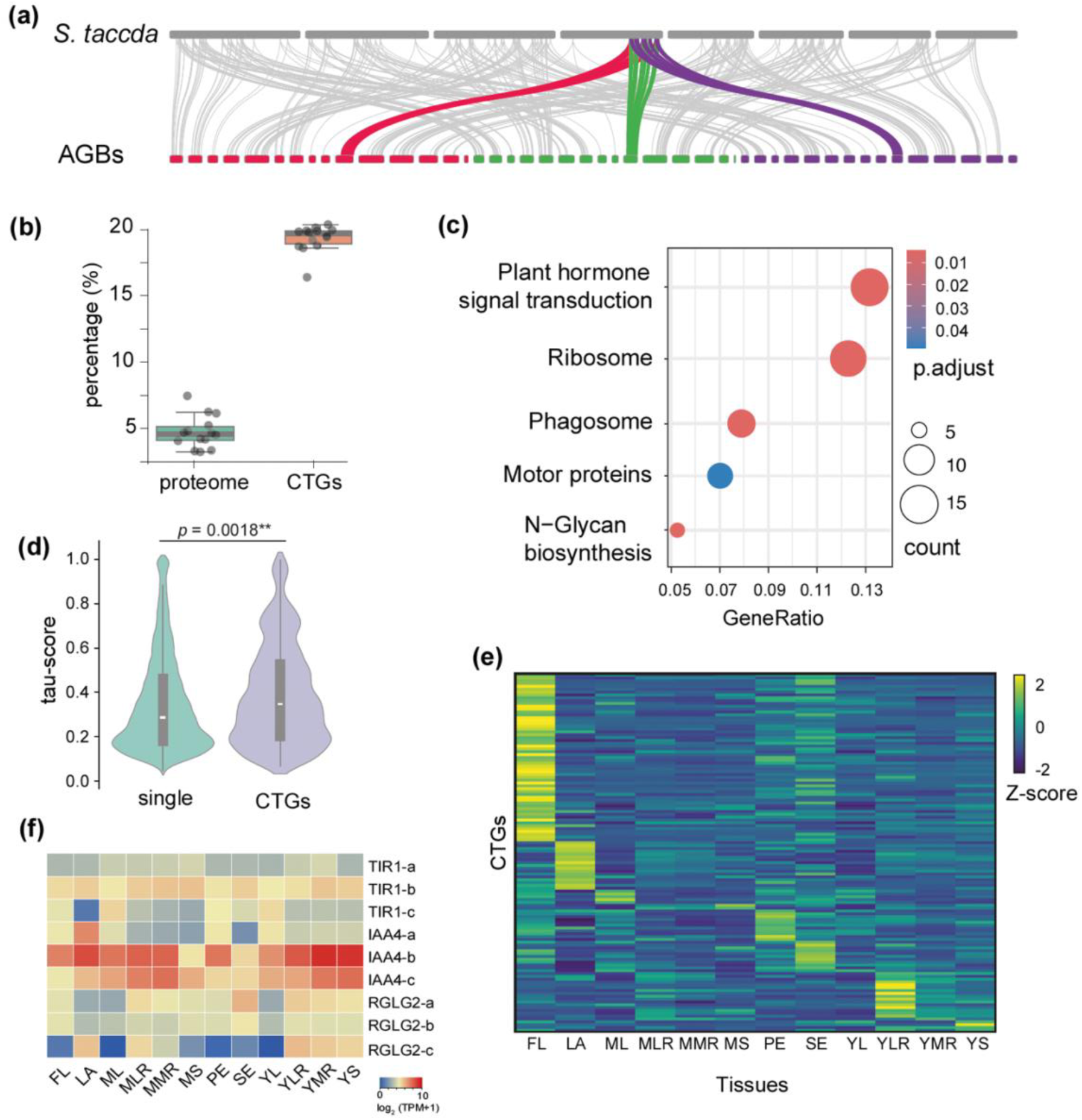
The conserved triplicated genes (CTGs). **a.** Distribution of CTGs on *S. taccada* and the pseudo genome and their syntenic relationships, with a hotspot of CTGs on the end of chromosome 4 of *S. taccada* highlighted; **b.** The percentage of transcription factors (TFs) in proteomes and CTGs in Asteraceae species to show TFs are significantly overrepresented in CTGs; **c.** Enriched KEGG pathways of the CTGs; **d.** Expression specificity (measured as *tau* score) of CTGs and single-copy genes in *T. kok-saghyz* tissues; **e.** The expression of CTGs in 12 different tissues of *T. kok-saghyz*. (FL-flower, LA-latex, ML-mature leaf, MLR-mature lateral root, MMR-mature main root, MS-mature stem, PE-peduncle, SE-seed, YL-young leaf, YLR-young lateral root, YMR-young mature root, YS-young stem. The expression was measured as TPM value and normalized as z-score; **f.** Expression of auxin related genes in various *T. kok-saghyz* tissues.

### CTGs are overrepresented by transcription factors and enriched in plant hormone pathways

Next, we ask what types of genes are the CTGs, and what is the evolutionary significance of their conservation? Initial inspection of the CTGs found that many of them are transcription factors (TFs), such as *MYB*, *WRKY* and *NAC* (Table S3). We then screened the proteomes of Asteraceae species for TFs (https://planttfdb.gao-lab.org/prediction.php, accessed at 26-05-2024), and identified 1,705-2,650 TFs, which on average make up 4.1% of their proteomes. However, on average 18.5% of the 471 CTGs are TFs, which is significant overrepresentation (Fig. 6b). Gene function enrichment analysis identified four significantly overrepresented KEGG pathways, with the top one being plant hormone signalling pathways, specifically the auxin and ethylene related biological processes (Fig. 6c). Further gene ontology (GO) enrichment analysis found that the CTGs are significantly enriched in 179 GO terms (p<0.05) which are clustered into several modules, including flower development and plant hormones (Fig. S12).

We then explored the expression specificity and divergence of the CTGs in various tissues of dandelion (30). We found that compared to single-copy genes, the CTGs are significantly specialized in expression pattern, namely the expression of CTGs has higher tissue specificity compared to single-copy genes (Fig. 6d). Interestingly, almost half of the CTGs (tau>0.5) are preferentially expressed in flowers (Fig. 6f). Consistently, there are 30 CTGs of which the Arabidopsis orthologs are known to be involved in flower development (Table S3), including the MADS domain transcription factor *SOC1* which is a central floral integrator (31). There are also several diverse auxin signalling related genes which show differentiated gene expression between the triplicated paralogs (Fig. 6e). This includes TIR1 (the auxin receptor whose function is to mark Aux/IAA’s for degradation in the presence of auxin), IAA4 (a repressor of the auxin response, and turned over by TIR1) and RGLG2 (a RING domain ubiquitin E3 ligase targeting auxin pathway proteins for degradation).

## Discussion

The genome triplication of the Asteraceae, held to have occurred via a two-step process, is widely accepted (12, 17, 20). However, the contribution of the progenitor genomes to modern Asteraceae genomes and the biological significance of the paleo-hexaploidization requires further research. Previous studies to decipher the genomic legacy of this event rely on comparisons between Asteraceae and the far-related *Vitis vinifera* (20) or are limited to pairwise comparisons of specific regions (24). Our multi-genome synteny analysis, situated within the context of the Asteraceae phylogeny, allows us to investigate the genomic conservation and/or diversification at multiple scales.

At the chromosome level, we observed that the progenitor genomes had evolved through extensive fissions, fusions, inversions and translocations, resulting in the modern Asteraceae genomes being complex mosaics, with variability evident between lineages. Such highly dynamic evolution is different from the recently sequenced paleo-hexaploid *Platanus* genome which has retained almost the same karyotype as its ancestor (32). Chromosomal rearrangements are hypothesized to facilitate adaptation and genomic innovation, both by bringing together previously unlinked adaptive alleles and by creating regions of low recombination that facilitate the linkage of adaptive alleles (33). In addition, we also detected extensive genomic exchange among progenitor genomes which further increase the genomic diversity of Asteraceae. At the gene level, we identified Asteraceae-wide and lineage-specific synteny clusters.

The evolutionary significance of gene synteny is not well understood, however, evidence for the co-regulation of neighbour genes is accumulating (34, 35). Colocalization of genes in genomes may facilitate preservation of favourable allele combinations between epistatic loci or coregulation of functionally related genes (36, 37). One of the most outstanding examples is the biosynthetic gene clusters which have been widely reported in plant specialized metabolisms (38–40). Additionally, we found there is no significant gene fractionation bias among progenitor genomes in Asteraceae, suggesting that they have contributed equally to the gene content of Asteraceae; This pattern is different from Brassicaceae ancient polyploidy events in which clear subgenome dominance has been observed (41). This may suggest that the ancient Asteraceae WGD events were autopolyploidization or allopolyploids with frequent homoeologous exchanges (42). However, further comparative genomic analysis involving the intermediate paleo-tetraploids, such as Calyceraceae, is needed to understand the nature of the two-step triplication. In short, we demonstrated that the two-step genome triplication and subsequent genomic exchanges, chromosomal shuffling and gene fractionation strongly reshaped the architecture and gene content of modern Asteraceae genomes.

We also highlight the CTGs identified in Asteraceae genomes. Different from previous work based on pairwise comparison (24), our comparative genome analysis includes multiple genomes from diverse lineages, therefore accounting for lineage-specific gene loss. We found that the CTGs are significantly overrepresented by TFs, mainly plant hormone related regulatory genes. A previous study on the retention of Asteraceae paleologs based on evolutionary distance (*K*s) of gene pairs found that transcription and other regulatory functions were significantly underrepresented (16). It is likely that the *K*s based paleologs include both triplicated and duplicated genes, and non-syntenic genes, representing an assemblage of more broad paralogs than the CTGs identified here. Our study offers a new perspective on the conservation and diversification of paleologs in Asteraceae. Together, these observations indicated that retention of the triplication derived paleologs in Asteraceae is heterogenous among function categories.

Understanding the genetic mechanisms for morphological innovation in Asteraceae, such as capitulum, is one of the main tasks of the Asteraceae community (43, 44). The origin of complex organ is believed to be primarily driven by rewiring of the existing regulatory networks (45), for example the origin of roots in vascular plants (46). It is suggested that the morphogenesis of the flower-like capitulum in Asteraceae is driven by recruitment of existing conserved developmental regulators (44). Intriguingly, genes annotated to flower development and plant hormones are significantly enriched among the CTGs; for example, *SOC1*, a MADS TF at the centre of flowering network (31), has retained its three paleologs in all representative Asteraceae subfamilies, likely being conserved in both copy number and synteny. Auxin plays essential role in the development of Asteraceae capitulum (43, 47, 48). Phytohormones such as gibberellin, jasmonate and abscisic acid are also important mediators in floral organ morphogenesis (49, 50); for example, jasmonate participates in the regulation of late-stage floral organ development (51). Given the widely occurred flower heteromorphosis in Asteraceae (52–54), sub-functionalization of the related CTG paleologs and their involvement in the determination of floret identity is a reasonable hypothesis. In addition, the regulation of gene expression by TFs is position dependent, namely activation or repression of gene transcription depends on the precise position of TF relative to the target gene (55). Understanding the conservation or diversification of gene synteny across lineages, for example the CTGs and their neighbour genes, can provide insights into the evolution of regulation networks, eventually the phenotypic diversity.

Advances in genome sequencing technology have precipitated comparative genomic research in Asteraceae (2), and initial breakthroughs have also been made regarding the genetic mechanisms underpinning key innovative traits, such as the capitulum (43, 56). Given the considerable variability in genome architecture between lineages and the high evolutionary dynamics of ALGs in Asteraceae, phylogenetic coverage is a crucial aspect for comparative analysis. However, the currently available Asteraceae genome data are biased towards certain taxonomic groups and are phylogenetically unevenly distributed, with only 10 of the 57 tribes represented and nine of them from subfamily Asteroideae. Early diverging lineages, such as Barnadesioideae and Famatinanthoideae, should be a priority for future sequencing projects. Nonetheless, our synteny-phylogenomic framework, the 15 ALGs, and a suite of tools to interpret whole genome synteny and regional syntenic clusters will accelerate the comparative genomic study in Asteraceae.

## Materials and Methods

### Genome data and quality assessment

As of the time of our analysis, a total of 61 assemblies from 35 distinct species were publicly available (https://www.ncbi.nlm.nih.gov/datasets/genomes/?taxon=4210), with most of them being generated in the past three years. As our main purpose was to build up a synteny-phylogenomic framework for Asteraceae, several criteria were applied to choose genomes for comparative analysis: 1) chromosome level assembly; 2) BUSCO scores; and OMArk scores; and 3) maximizing phylogenetic representation. This resulted in 20 Asteraceae genomes from 10 tribes in three subfamilies (Table S1, Fig. S1). In addition, *Scaevola taccada* from Goodeniaceae was included as outgroup for the comparative analysis. BUSCO analyses were performed using the eudicot single-copy genes (*eudicots_odb10*) as reference and conducted in BUSCO v5.8.2 (21) with default settings. As our analyses largely rely on the quality of the annotated protein coding regions (proteomes), we further assessed the quality of the proteomes using OMArk v0.3.0 (22) which places query protein sequences into known gene families and compares them to the expected families of the species’ lineage. Therefore, OMArk assesses not only the completeness but also the consistency of the gene repertoire as a whole relative to closely related species and reports likely contamination events. OMArk analyses were carried out with asterids as the reference lineage and the precomputed asterids orthologous groups was downloaded from OMA (https://omabrowser.org/oma/home/; accessed at 05-26-2023).

### Multi-genome macrosynteny analysis

Global synteny across the genome was inferred using the recently developed phylogeny-aware genome synteny tool GENESPACE (27). GENESPACE utilizes OrthoFinder (57) to first infer hierarchical ortholog groups, based on which the homologous genomic regions between genomes, namely syntenic blocks, were inferred. Therefore, GENESPACE is capable of distinguishing paralogous blocks and orthologous blocks. With the prior knowledge on the history of genome duplication in Asteraceae and ploidy status of Asteraceae species (Table S1), we carried out three different analyses to fulfil different objectives. Analysis 1: to construct genome synteny between the outgroup *S. taccada* and Asteraceae species, the hierarchical ortholog groups (hOGs) at node N1 (Fig. S1) were used, and the ploidy level for *S. taccada* was set as ‘1’ and ploidy level for Asteraceae species were set as ‘3’ for diploids and ‘3x’ for extant polyploids or extant diploids that have additional WGD events (Table S1), where x is ploidy level or counts of additional WGD after ancient triplication. For example, sunflower is an extant diploid but has undergone one WGD after the ancient triplication, so the ploidy level is ‘6’. Analysis 2: as we are specifically interested in the orthologous synteny between the outgroup S. taccada and Asteraceae species, we used the same hOGs as in analysis 1 while set the ploidy level as ‘1’ for all species. This identified the same orthologous synteny as in analysis 1 while excluded the paralogous synteny among Asteraceae species. Analysis 3: to construct the synteny among Asteraceae species, the hOGs at node N2 were used, and ploidy level for all diploids are ‘1’, and ‘1x’ for others as mentioned above. For all analysis, protein sequences with less than 30 amino acids, and hOGs scattered on >= 8*x locations (where x is the ploidy level for a given species) were included in OrthoFinder analysis but excluded in synteny inference. The macro-synteny across species was visualized as a riparian plot. Syntenic genes across species was obtained using the *pangene* function as implemented in GENESPACE and used for further analysis.

### Multi-genome microsynteny analysis

Microsynteny refers to the local conservation of orthologous gene order in genomic regions. We obtained the orthologous groups from the macrosynteny analysis using the *query_pangene* function as embedded in GENESPACE v1.3 (27). The syntenic orthologous genes across Asteraceae and Goodeniaceae genomes that are identified by GENESPACE were extracted and syntenic pairs were generated using custom scripts. Syntenic gene pairs were then used as inputs for *syntenet* (58) to perform phylogenetic profiling and to call microsynteny clusters. A microsynteny cluster refers to a homologous group in which the gene members are syntenic across genomes/species (23). The microsynteny of Asteraceae and Goodeniaceae genes was represented by heatmaps (Fig. 1b, Fig. S9) and networks (Fig. 1d). In addition, we also quantified the synteny gene clusters in different phylogenetic branches.

### Chromosome rearrangement estimates

*Scaevola taccada* was used as outgroup to infer the chromosome rearrangements in Asteraceae diploid species (Table S1). *S. taccada* is a diploid species and has no genome duplication after its divergence from Asteraceae-Calyceraceae lineage. Comparisons between *S. taccada* and the diploid Asteraceae species can infer the diploidization process (e.g. chromosome reversals, fusions, fissions, translocations) after the ancient genome triplication in Asteraceae. To infer the genomic rearrangements, the syntenic genes identified in microsynteny analysis were used as anchors to reconstruct synteny blocks. We used DRIMM-Synteny (26) (*cycleLengthThreshold=20*, *dustLengthThreshold=5*) to reconstruct synteny blocks shared by *S. taccada* and Asteraceae species, and blocks with at least 5 anchor genes were then used to infer rearrangement events using IAGS (59).

### Reconstruction of Ancestral linkage group

Genome rearrangement analysis revealed that the *A. lappa* genome exhibits the least rearrangement among the genomes analyzed in this study. In addition, *A. lappa* genome assembly is one of the best in both completeness and continuity (Table S1). Consequently, we used *A. lappa* to reconstruct ancestral linkage groups (ALGs) in the context of the ancient Asteraceae genome triplication. Initially, we employed DRIMM-Synteny to identify synteny segments between *A. lappa* and *S. taccada*, based on a syntenic depth ratio of 1:3. The synteny segments representing the *A. lappa* genome were visualized through chromosome bar painting using chromoMap v4.1.1 (60). Adjacent synteny segments (each containing ≥3 anchor genes) on *A. lappa* chromosomes were chained into syntenic blocks if their orthologous counterparts in the *S. taccada* genome were physically proximate. This approach assumes that such synteny segments share a common ancestry and belong to the same linkage group. As a result, 98 preliminary blocks were identified (Table S4), covering 90.15% of the *A. lappa* genome in terms of protein-coding gene counts.

To refine these blocks, we generated a genomic dot plot between the preliminary blocks and the *S. taccada* genome using MCScanX (61) (Fig. S4). Guided by the dot plot, the preliminary blocks were assigned to 15 × 3 blocks based on the principles of proximity and complementarity (Fig. 2b). These 15 × 3 blocks are presumed to represent the three subgenomes derived from the ancient Asteraceae genome polyploidization. Since these blocks were reconstructed using the extant *A. lappa* genome, it is likely that lineage-specific rearrangements have altered the chromosome architecture compared to the ancestral state.

To approximate the ancestral configuration, we used *S. taccada* as a reference and rearranged the preliminary blocks in both orientation and order to achieve a high degree of collinearity with *S. taccada* chromosomes (Fig. S7). The arrangements were accomplished using a custom developed pipeline (https://github.com/xiaoyezao/Asteraceae-synteny-phylogenomics) which utilized the functions from Pybedtools v0.9.1 (62) and RagTag v2.1.0 (63). Based on the redefined 15 × 3 blocks, we generated a pseudo genome.

### Subgenome phasing by phylogenetic analysis and Ks analysis

To assign the paralogous genomic blocks (namely the three sister blocks in each set of ALG) to biological subgenomes, we carried out phylogenomic analysis on the pseudo subgenomes with *S. taccada* as outgroup. The phylogenetic analysis was conducted for each set of the 15 ALGs using the syntenic genes. Because of gene fractionation, the syntenic genes may have been lost in one or more of the pseudo subgenomes. To overcome this, we performed sliding-window phylogenetic analyses. Windows of coding sequence (size = 5 genes, step = 5 genes) were generated for each ALG, and the windows with at least two genes in any subgenomes and *S. taccada* were used for phylogenetic inference. A concatenated protein sequence alignment for each window was generated using MAFFT v7.520 (*--genafpair; --maxiterate 1,000*) (64), and was cleaned using trimAl v1.4.1 (*-gt 0.6; -st 0.001*) (65). The best-fit amino acid substitution model was identified using ModelFinder (66), and the best maximum likelihood tree was inferred using RAxML v2.3.6 (67) with 200 bootstrapping replicates. The inconsistences between topologies were visualized in DensiTree (68).

To characterize the sequence divergence between the pseudo subgenomes, we carried out Ks (synonymous substitutions per site) analysis. The syntenic gene pairs of subgenomes were used to calculate ingroup Ks values, and the syntenic gene pairs between the subgenomes and *S. taccada* genome were used to calculate ingroup-outgroup Ks values. The calculations were conducted for each ALGs independently using the *ksd* function in wgd2 v2.0.38 (69) with default settings. Distributions of Ks were visualized by histogram and violin plotting and the significance of the divergence between (sub)genome pairs was tested using Tukey HSD for multiple comparison of means.

### Gene fractionation analyses

Gene fractionation pattern in the three subgenomes was analysed by comparing the ALGs with *S. taccada* genome. The reconstructed ALGs were aligned with the outgroup *S. taccada* genome as described in macrosynteny analysis. The syntenic genes were obtained using the *pangene* function in GENESPACE with *S. taccada* genome as reference. The fractionation of genes in ALGs were characterized by calculating the retention of syntenic genes on ALGs per 100-gene window along the reference chromosomes.

### Frackify analyses

To further validate fractionation patterns, we ran the Frackify pipeline, which identifies paleo-paralogous gene pairs from interspecific and intraspecific syntenic comparisons (29). We used default settings for MCScanX (61) to make syntenic comparisons in Asteraceae species and *Scaevola taccada* as an outgroup for each analysis. For each collinear gene pair, we calculated their KS using the *add_ka_and_ks_to_collinearity.pl* script available through the MCScanX repository. We then identified peaks in the KS distributions using the *find_peaks*, *nparam_density*, and *gaussian_kde* functions from the Numpy and Scipy python libraries (70, 71). These data cumulatively were used as input for Frackify (29).

### Identification of conserved triplicated genes (CTGs)

To identify conserved triplicated genes, a multi-species gene fractionation analysis was conducted. In contrast to the pairwise approach, which entails the comparison of a single Asteraceae species with an outgroup species on each occasion, the multi-species analysis allows for the examination of the presence of a specific gene with all three paleo paralogs in multiple species simultaneously. To perform the analysis, the proteomes of the species used in this study were subjected to phylogenetic profiling in OrthoFinder v2.5.5 (57) with a predefined phylogenetic topology. The hierarchical orthologous groups at node 1 (see Fig. S1) were employed to identify the paleo paralogs through the utilisation of a synteny analysis approach, as delineated in the microsynteny analysis section. The most rigorous criterion would be to require the presence of all three paleo paralogs in all species. However, to account for the incompleteness of the genome assembly and annotation (although most genomes used here have a BUSCO score of >90%), two different, relaxed criteria were employed to identify the CTGs. Firstly, a first cutoff was applied, whereby for a given gene, the three paleo paralogs were required to be present in >10 species in total of 14. Secondly, a second cutoff was employed, whereby the three paleo paralogs were required to be present in >75% of species within each of the three Asteraceae subfamilies. The intersection set from the two analyses were then taken as the CTGs.

### Gene expression analyses

Gene expression data from previous studies (30) which includes diverse tissues and developmental stages was used. RNAseq reads were mapped to the reference genome using STAR v2.7.11b (72), and TPM (transcripts per million) values were calculated by TPMcalculator v0.0.5 (73). Genes that are expressed (TPM >= 5) in at least one tissue are used for further analysis. Gene expression specificity was estimated used an extended *tau* score algorithm (74).

### GO and KEGG enrichment analyses

Overrepresented GO (Gene Ontology) and KEGG (Kyoto Encyclopedia of Genes and Genomes) terms in target gene sets were identified using Fisher’s exact test and the significance of the test (adjusted p value) were estimated using Benjamini-Hochberg method (75) by accounting for multiple tests. To perform the function enrichment analyses, we used lettuce (*Lactuca sativa*) as the reference. The protein coding genes were first functionally annotated using eggNOGMapper (http://eggnog-mapper.embl.de/), and then a custom lettuce-specific database that links gene ID with GO and KEGG terms was built using custom script: 1_build_OrgDB.R (https://github.com/xiaoyezao/Alpine-Plant-Genomics). The overrepresented GO and KEGG terms in gene sets of interests, either the conserved syntenic genes or the CTGs were explored using clusterProfiler v4.0 (76) as pipelined in custom script: 2_GO_KEGG.R (https://github.com/xiaoyezao/Alpine-Plant-Genomics). GO and KEGG enrichment analysis were performed using *‘enricher’* (pvalueCutoff = 0.05, qvalueCutoff = 1) in the R package “clusterProfiler”. The overrepresented GO and KEGG terms were visualized by bar plot and overlapped or related terms were clustered by *‘pairwise_termsim’* function in clusterProfiler.

## Acknowledgments

This publication is part of the LettuceKnow project (with project number 1.1 of the research Perspective Program P19-17 which is (partly) financed by the Dutch Research Council (NOW; TTW) and the breeding companies BASF, Bejo Zaden B.V., Limagrain, Enza Zaden Research & Development B.V., Rijk Zwaan Breeding B.V., Syngenta Seeds B.V., and Takii and Company Ltd. John Lovell is supported by the U.S. Department of Energy Joint Genome Institute (https://ror.org/04xm1d337), a DOE Office of Science User Facility, is supported by the Office of Science of the U.S. Department of Energy operated under Contract No. DE-AC02-05CH11231.

## Author Contributions

M.E.S. and T.F. conceived and designed the study; T.F. collected the data and performed the analyses with help from J.L.; M.M. performed the Frackify analysis; T.F. and M.E.S. wrote the paper with contributions from M.M., M.B., L.H.R., R.M. and J.L.; All authors read and approved the final manuscript.

## Competing Interest Statement

The authors declare no competing interest.

## Classification

Biological Sciences, Evolution

## Notes

### Competing Interest Statement

The authors have declared no competing interest.

## References

1. R. Pozner, C. Zanotti, L. A. Johnson, Evolutionary origin of the Asteraceae capitulum: Insights from Calyceraceae. Am J Bot 99, 1–13 (2012).

2. L. Palazzesi, et al., Asteraceae as a model system for evolutionary studies: from fossils to genomes. Botanical Journal of the Linnean Society 200, 143–164 (2022).

3. H. J. B. Birks, High-elevation limits and the ecology of high-elevation vascular plants: legacies from Alexander von Humboldt. Front Biogeogr 13, 1–18 (2021).

4. D. Vitales, P. Fernández, T. Garnatje, S. Garcia, Progress in the study of genome size evolution in Asteraceae: analysis of the last update. Database 2019, 1–13 (2019).

5. J. R. Mandel, et al., A fully resolved backbone phylogeny reveals numerous dispersals and explosive diversifications throughout the history of Asteraceae. Proc Natl Acad Sci U S A 116, 14083–14088 (2019).

6. H. Badouin, et al., The sunflower genome provides insights into oil metabolism, flowering and Asterid evolution. Nature 546, 148–152 (2017).

7. D. Scaglione, et al., The genome sequence of the outbreeding globe artichoke constructed de novo incorporating a phase-aware low-pass sequencing strategy of F1 progeny. Sci Rep 6, 19427 (2016).

8. S. Reyes-Chin-Wo, et al., Genome assembly with in vitro proximity ligation data and whole-genome triplication in lettuce. Nat Commun 8, 14953 (2017).

9. J. L. Panero, et al., Resolution of deep nodes yields an improved backbone phylogeny and a new basal lineage to study early evolution of Asteraceae. Mol Phylogenet Evol 80, 43–53 (2014).

10. V. D. Barreda, et al., Early evolution of the angiosperm clade Asteraceae in the Cretaceous of Antarctica. Proceedings of the National Academy of Sciences 112, 10989–10994 (2015).

11. J. R. Mandel, et al., A Target Enrichment Method for Gathering Phylogenetic Information from Hundreds of Loci: An Example from the Compositae. Appl Plant Sci 2, 1300085 (2014).

12. C. H. Huang, et al., Multiple Polyploidization Events across Asteraceae with Two Nested Events in the Early History Revealed by Nuclear Phylogenomics. Mol Biol Evol 33, 2820–2835 (2016).

13. C. Zhang, et al., Phylotranscriptomic insights into Asteraceae diversity, polyploidy, and morphological innovation. J Integr Plant Biol 63, 1273–1293 (2021).

14. A. Susanna, et al., The classification of the Compositae: A tribute to Vicki Ann Funk (1947– 2019). Taxon 69, 807–814 (2020).

15. A. R. Zuntini, et al., Phylogenomics and the rise of the angiosperms. Nature 629, 843–850 (2024).

16. M. S. Barker, et al., Multiple paleopolyploidizations during the evolution of the Compositae reveal parallel patterns of duplicate gene retention after millions of years. Mol Biol Evol 25, 2445–2455 (2008).

17. M. S. Barker, et al., Most compositae (Asteraceae) are descendants of a paleohexaploid and all share a paleotetraploid ancestor with the calyceraceae. Am J Bot 103, 1203–1211 (2016).

18. B. Liu, et al., Mikania micrantha genome provides insights into the molecular mechanism of rapid growth. Nat Commun 11, 340 (2020).

19. W. Fan, et al., The genomes of chicory, endive, great burdock and yacon provide insights into Asteraceae palaeo-polyploidization history and plant inulin production. Mol Ecol Resour (2022). 10.1111/1755-0998.13675.

20. X. Kong, et al., Two-step model of paleohexaploidy, ancestral genome reshuffling and plasticity of heat shock response in Asteraceae. Hortic Res 10 (2023).

21. M. Seppey, M. Manni, E. M. Zdobnov, “BUSCO: Assessing genome assembly and annotation completeness” in Methods in Molecular Biology, (2019), pp. 227–245.

22. Y. Nevers, et al., Quality assessment of gene repertoire annotations with OMArk. Nat Biotechnol 1–17 (2024). 10.1038/s41587-024-02147-w.

23. T. Zhao, M. E. Schranz, Network-based microsynteny analysis identifies major differences and genomic outliers in mammalian and angiosperm genomes. Proceedings of the National Academy of Sciences 116, 2165–2174 (2019).

24. F. Shen, et al., Comparative genomics reveals a unique nitrogen-carbon balance system in Asteraceae. Nat Commun 14, 1–14 (2023).

25. S. Ou, J. Chen, N. Jiang, Assessing genome assembly quality using the LTR Assembly Index (LAI). Nucleic Acids Res 46, e126 (2018).

26. S. K. Pham, P. A. Pevzner, DRIMM-Synteny: decomposing genomes into evolutionary conserved segments. Bioinformatics 26, 2509–2516 (2010).

27. J. T. Lovell, et al., GENESPACE tracks regions of interest and gene copy number variation across multiple genomes. Elife 11, 1–20 (2022).

28. K. Do Kim, et al., Chromosome-level genome assembly of milk thistle (Silybum marianum (L.) Gaertn.). Sci Data 11, 342 (2024).

29. M. T. W. McKibben, M. S. Barker, “Applying Machine Learning to Classify the Origins of Gene Duplications” in Methods in Molecular Biology, (2023), pp. 91–119.

30. T. Lin, et al., Extensive sequence divergence between the reference genomes of Taraxacum kok-saghyz and Taraxacum mongolicum. Sci China Life Sci 65, 515–528 (2022).

31. R. G. H. Immink, et al., Characterization of SOC1’s Central Role in Flowering by the Identification of Its Upstream and Downstream Regulators. Plant Physiol 160, 433–449 (2012).

32. X. Yan, et al., Genome evolution of the ancient hexaploid Platanus × acerifolia (London planetree). Proceedings of the National Academy of Sciences 121, 2017 (2024).

33. Z. Liu, et al., Chromosomal Fusions Facilitate Adaptation to Divergent Environments in Threespine Stickleback. Mol Biol Evol 39 (2022).

34. P. Fan, et al., Evolution of a plant gene cluster in Solanaceae and emergence of metabolic diversity. Elife 9, 1–26 (2020).

35. S. Wu, et al., Convergent gene clusters underpin hyperforin biosynthesis in St John’s wort. New Phytologist 235, 646–661 (2022).

36. T. Makino, A. McLysaght, Positionally biased gene loss after whole genome duplication: Evidence from human, yeast, and plant. Genome Res 22, 2427–2435 (2012).

37. T. Makino, A. McLysaght, Interacting Gene Clusters and the Evolution of the Vertebrate Immune System. Mol Biol Evol 25, 1855–1862 (2008).

38. L. Mao, et al., Genomic evidence for convergent evolution of gene clusters for momilactone biosynthesis in land plants. Proc Natl Acad Sci U S A 117, 12472–12480 (2020).

39. H. Fang, et al., A monocot-specific hydroxycinnamoylputrescine gene cluster contributes to immunity and cell death in rice. Sci Bull (Beijing) 66, 2381–2393 (2021).

40. G. Polturak, A. Osbourn, Defense-related phenylpropanoid biosynthetic gene clusters in rice. Sci Bull (Beijing) 67, 13–16 (2022).

41. N. V. Hoang, et al., The Gynandropsis gynandra genome provides insights into whole-genome duplications and the evolution of C4 photosynthesis in Cleomaceae. Plant Cell 35, 1334–1359 (2023).

42. A. S. Mason, J. F. Wendel, Homoeologous Exchanges, Segmental Allopolyploidy, and Polyploid Genome Evolution. Front Genet 11, 1014 (2020).

43. T. Zhang, P. Elomaa, Development and evolution of the Asteraceae capitulum. New Phytologist 242, 33–48 (2024).

44. P. Elomaa, Y. Zhao, T. Zhang, Flower heads in Asteraceae—recruitment of conserved developmental regulators to control the flower-like inflorescence architecture. Hortic Res 5, 36 (2018).

45. S. B. Carroll, Evo-Devo and an Expanding Evolutionary Synthesis: A Genetic Theory of Morphological Evolution. Cell 134, 25–36 (2008).

46. W. Liu, L. Xu, Recruitment of IC-WOX Genes in Root Evolution. Trends Plant Sci 23, 490–496 (2018).

47. N. Zoulias, S. H. C. Duttke, H. Garcês, V. Spencer, M. Kim, The Role of Auxin in the Pattern Formation of the Asteraceae Flower Head (Capitulum). Plant Physiol 179, 391–401 (2019).

48. T. Zhang, et al., Phyllotactic patterning of gerbera flower heads. Proc Natl Acad Sci U S A 118, 1–11 (2021).

49. E. R. Lampugnani, A. Kilinc, D. R. Smyth, Auxin controls petal initiation in Arabidopsis. Development 140, 185–194 (2013).

50. G. Ren, et al., GhWIP2, a WIP zinc finger protein, suppresses cell expansion in Gerbera hybrida by mediating crosstalk between gibberellin, abscisic acid, and auxin. New Phytologist 219, 728–742 (2018).

51. Y. Cheng, X. Dai, Y. Zhao, Auxin biosynthesis by the YUCCA flavin monooxygenases controls the formation of floral organs and vascular tissues in Arabidopsis. Genes Dev 20, 1790–1799 (2006).

52. T. Zhang, P. Elmonaa, Understanding capitulum development: Gerbera hybrida inflorescence meristem as an experimental system. The International Compositae Alliance 1, 53 (2022).

53. N. G. Bergh, G. A. Verboom, Anomalous capitulum structure and monoecy may confer flexibility in sex allocation and life history evolution in the Ifloga lineage of paper daisies (Compositae: Gnaphalieae). Am J Bot 98, 1113–1127 (2011).

54. T. Zhang, P. Elomaa, Don’t be fooled: false flowers in Asteraceae. Curr Opin Plant Biol 59, 101972 (2021).

55. S. H. Duttke, et al., Position-dependent function of human sequence-specific transcription factors. Nature 631, 891–898 (2024).

56. V. Gurung, S. Muñoz-Gómez, D. S. Jones, Putting heads together: Developmental genetics of the Asteraceae capitulum. Curr Opin Plant Biol 81, 102589 (2024).

57. D. M. Emms, S. Kelly, OrthoFinder: Phylogenetic orthology inference for comparative genomics. Genome Biol 20, 1–14 (2019).

58. F. Almeida-Silva, T. Zhao, K. K. Ullrich, M. E. Schranz, Y. Van de Peer, syntenet: an R/Bioconductor package for the inference and analysis of synteny networks. Bioinformatics 39 (2023).

59. S. Gao, et al., IAGS: Inferring Ancestor Genome Structure under a Wide Range of Evolutionary Scenarios. Mol Biol Evol 39 (2022).

60. L. Anand, C. M. Rodriguez Lopez, ChromoMap: an R package for interactive visualization of multi-omics data and annotation of chromosomes. BMC Bioinformatics 23, 33 (2022).

61. Y. Wang, et al., MCScanX: a toolkit for detection and evolutionary analysis of gene synteny and collinearity. Nucleic Acids Res 40, e49–e49 (2012).

62. R. K. Dale, B. S. Pedersen, A. R. Quinlan, Pybedtools: a flexible Python library for manipulating genomic datasets and annotations. Bioinformatics 27, 3423–3424 (2011).

63. M. Alonge, et al., Automated assembly scaffolding using RagTag elevates a new tomato system for high-throughput genome editing. Genome Biol 23, 258 (2022).

64. K. Katoh, D. M. Standley, MAFFT multiple sequence alignment software version 7: Improvements in performance and usability. Mol Biol Evol 30, 772–780 (2013).

65. S. Capella-Gutiérrez, J. M. Silla-Martínez, T. Gabaldón, trimAl: A tool for automated alignment trimming in large-scale phylogenetic analyses. Bioinformatics 25, 1972–1973 (2009).

66. S. Kalyaanamoorthy, B. Q. Minh, T. K. F. Wong, A. von Haeseler, L. S. Jermiin, ModelFinder: fast model selection for accurate phylogenetic estimates. Nat Methods 14, 587–589 (2017).

67. A. M. Kozlov, D. Darriba, T. Flouri, B. Morel, A. Stamatakis, RAxML-NG: a fast, scalable and user-friendly tool for maximum likelihood phylogenetic inference. Bioinformatics 35, 4453– 4455 (2019).

68. R. R. Bouckaert, DensiTree: making sense of sets of phylogenetic trees. Bioinformatics 26, 1372–1373 (2010).

69. H. Chen, A. Zwaenepoel, Y. Van de Peer, wgd v2: a suite of tools to uncover and date ancient polyploidy and whole-genome duplication. Bioinformatics 40 (2024).

70. C. R. Harris, et al., Array programming with NumPy. Nature 585, 357–362 (2020).

71. P. Virtanen, et al., SciPy 1.0: fundamental algorithms for scientific computing in Python. Nat Methods 17, 261–272 (2020).

72. A. Dobin, et al., STAR: Ultrafast universal RNA-seq aligner. Bioinformatics 29, 15–21 (2013).

73. R. V. Alvarez, L. S. Pongor, L. Mariño-Ramírez, D. Landsman, TPMCalculator: One-step software to quantify mRNA abundance of genomic features. Bioinformatics 35, 1960–1962 (2019).

74. H. B. Lüleci, A. Yılmaz, Robust and rigorous identification of tissue-specific genes by statistically extending tau score. BioData Min 15, 31 (2022).

75. Y. Benjamini, Y. Hochberg, On the adaptive control of the false discovery rate in multiple testing with independent statistics. Journal of Educational and Behavioral Statistics 25, 60–83 (2000).

76. T. Wu, et al., clusterProfiler 4.0: A universal enrichment tool for interpreting omics data. The Innovation 2, 100141 (2021).

