## Supplementary figures and images for "Phylogenomic synteny analysis tracks conserved ancient polyploid-derived triplicated genomic blocks across Asteraceae genomes"

### Fig. S1

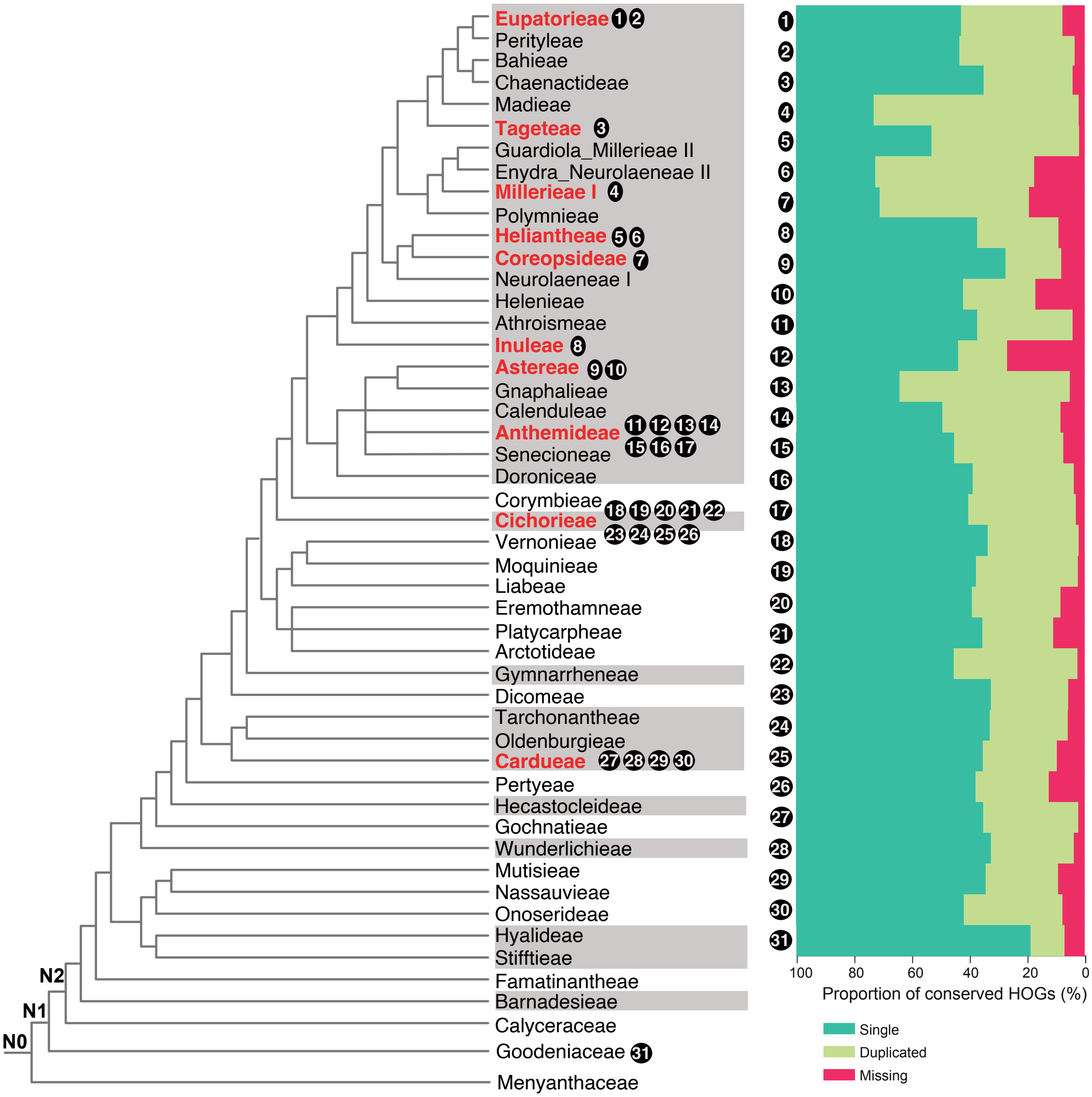

### Fig. S2

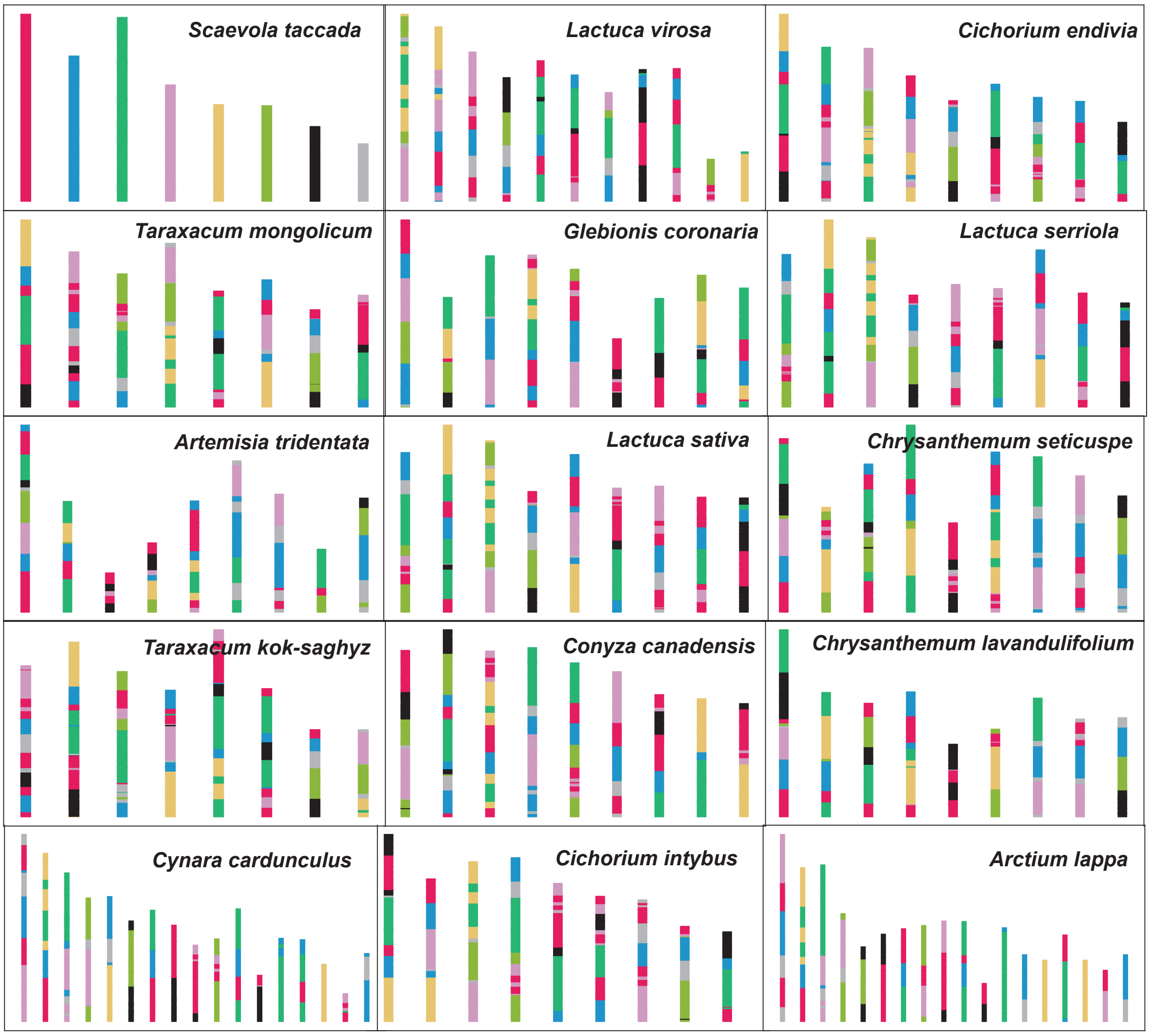

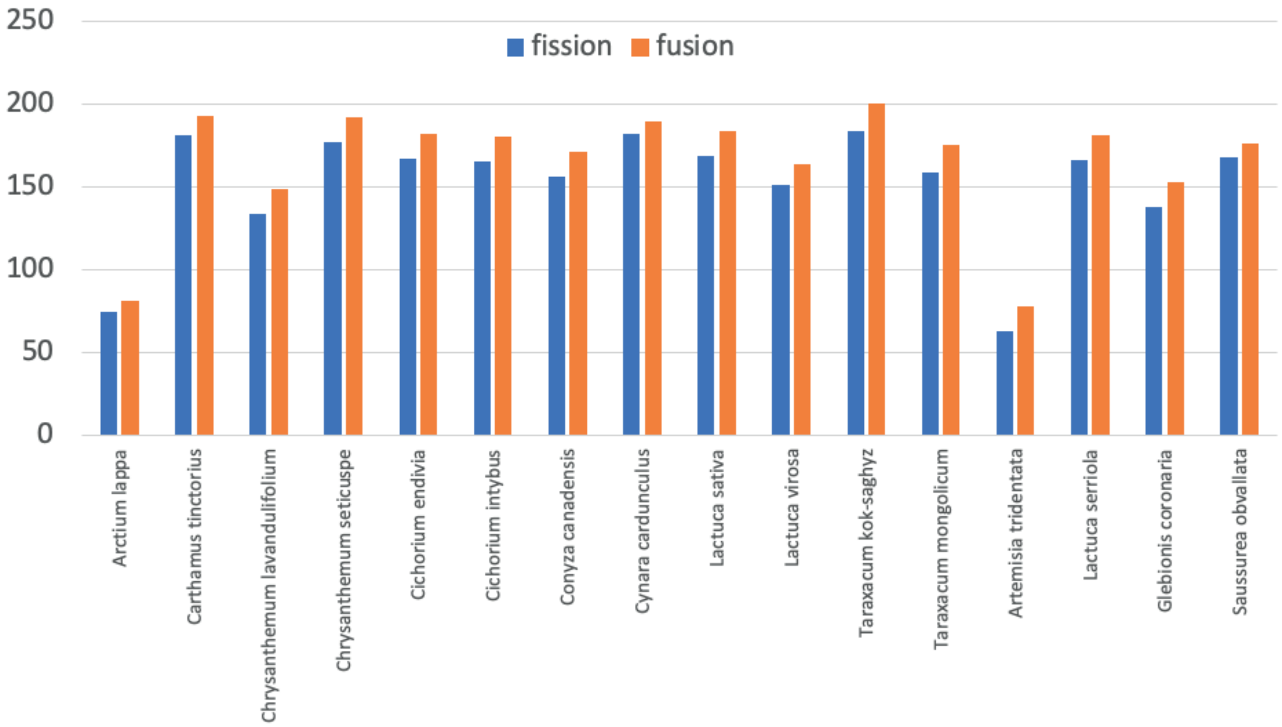

### Fig. S3

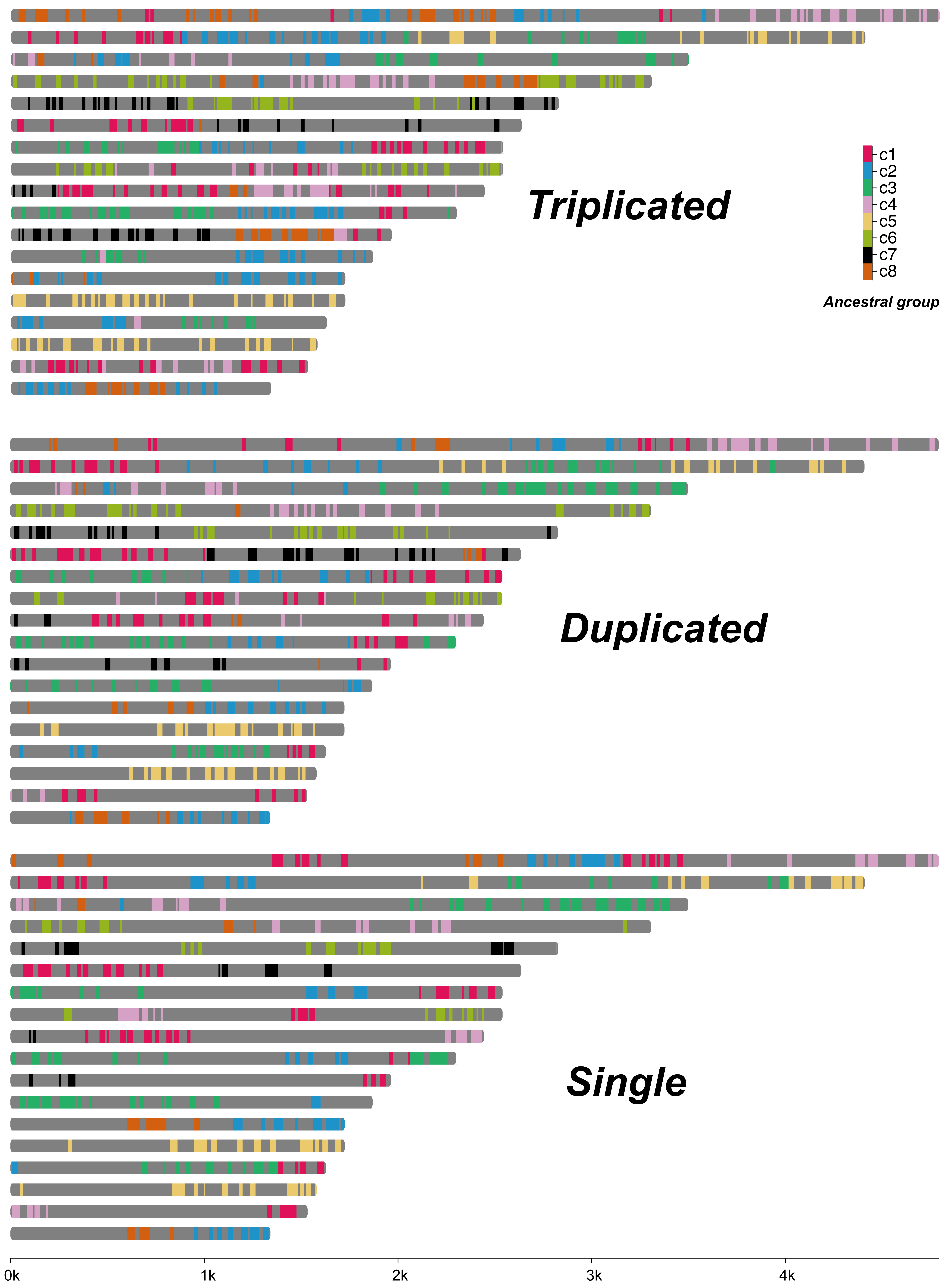

### Fig. S4

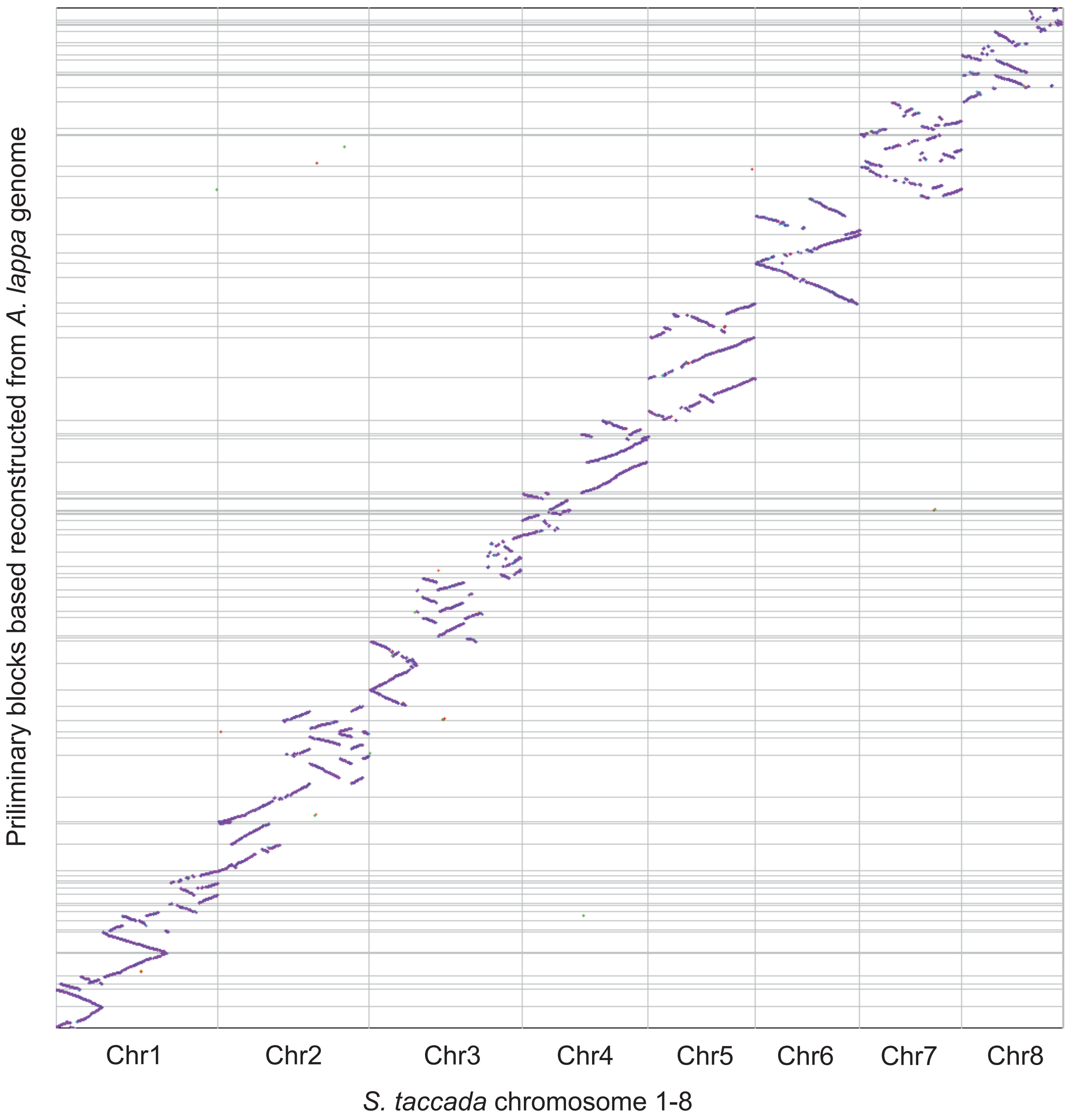

### Fig. S5

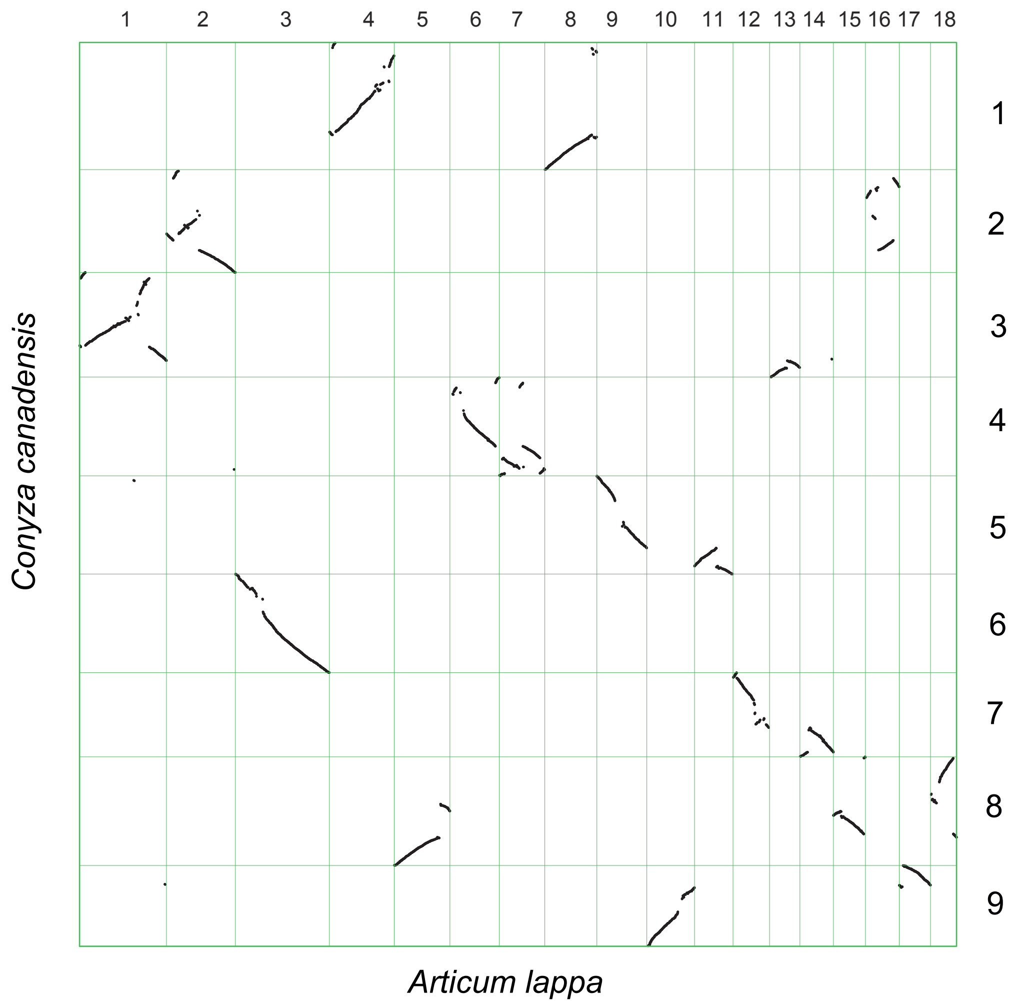

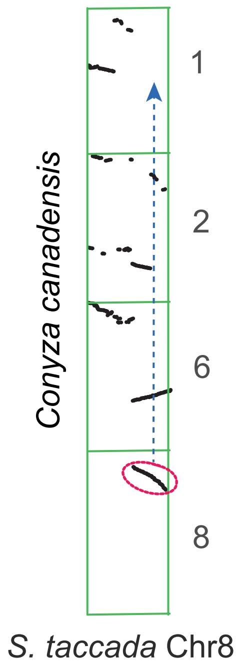

### Fig. S6

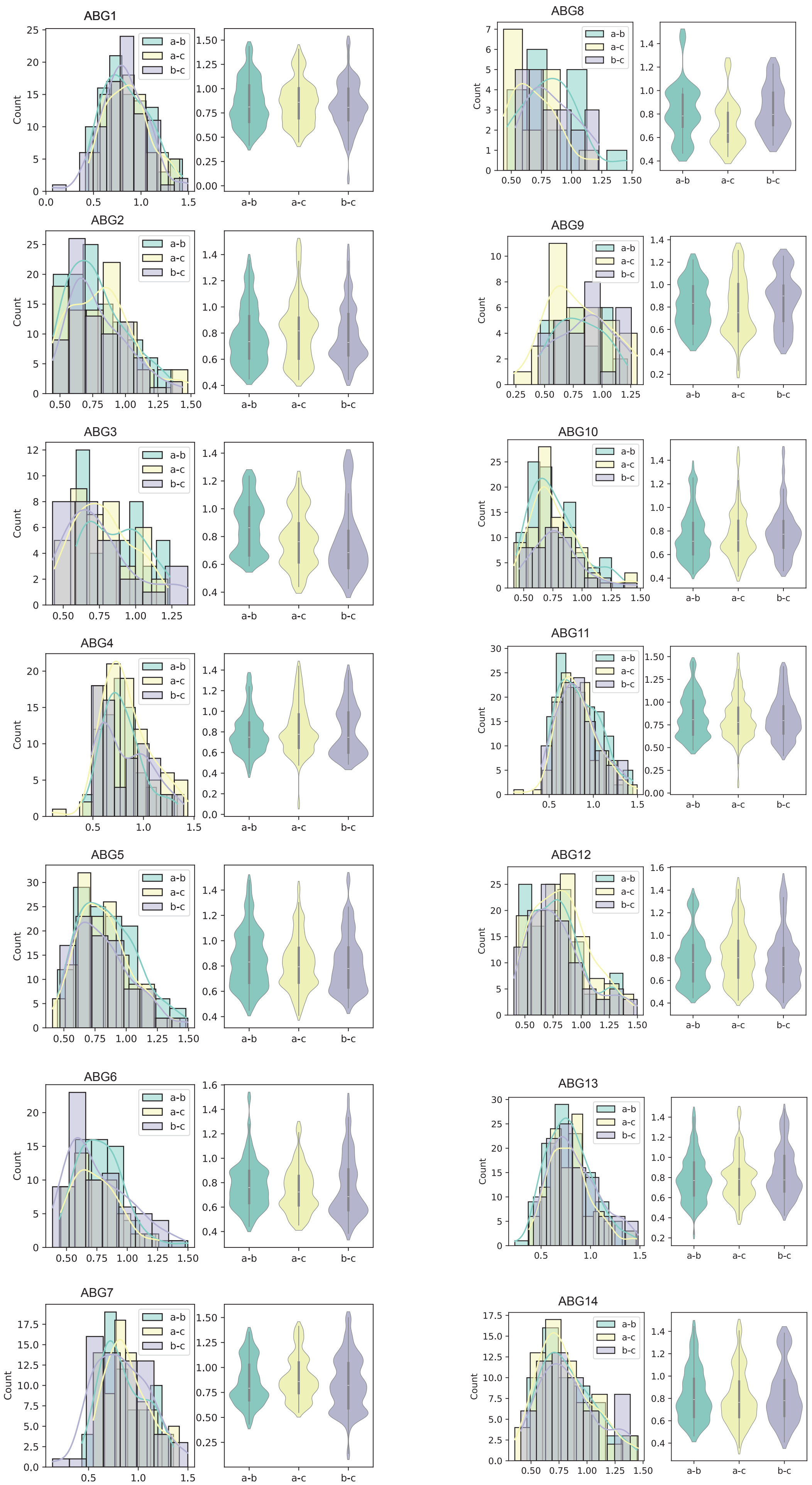

### Fig. S7

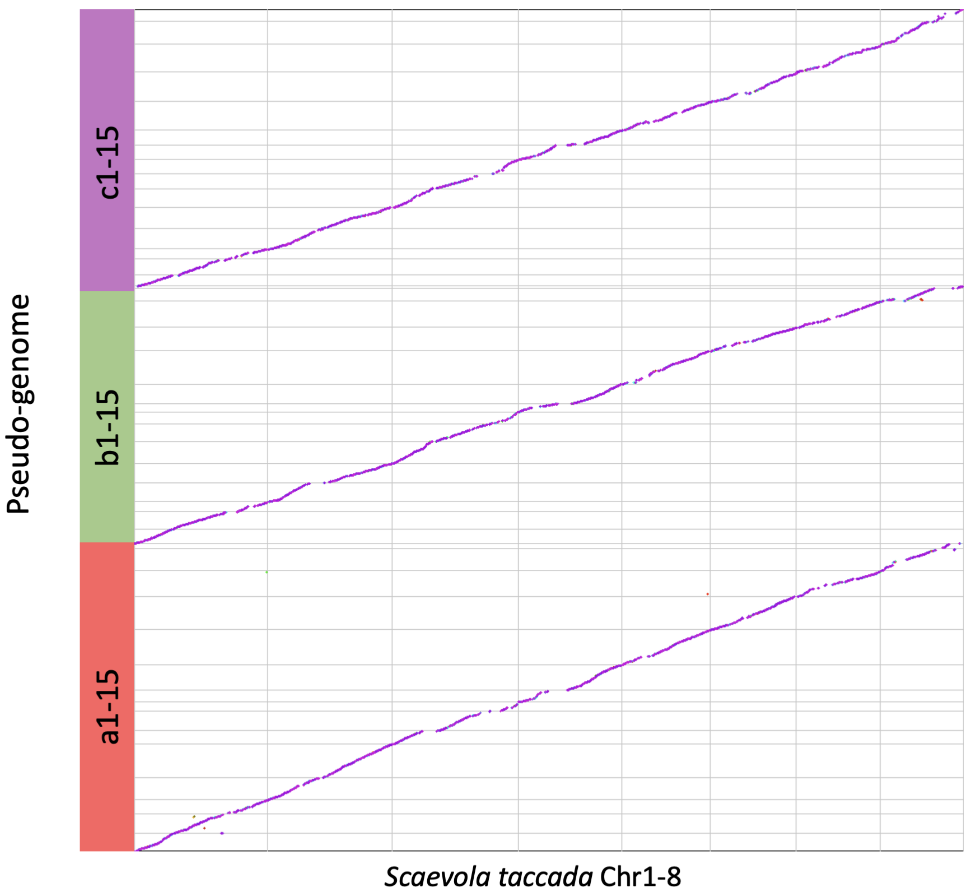

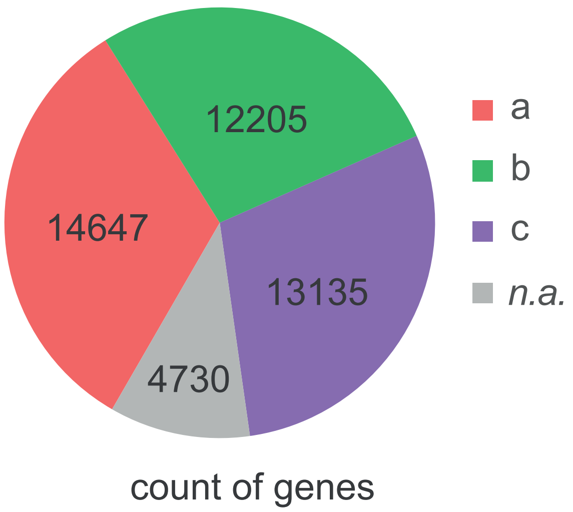

### Fig. S8

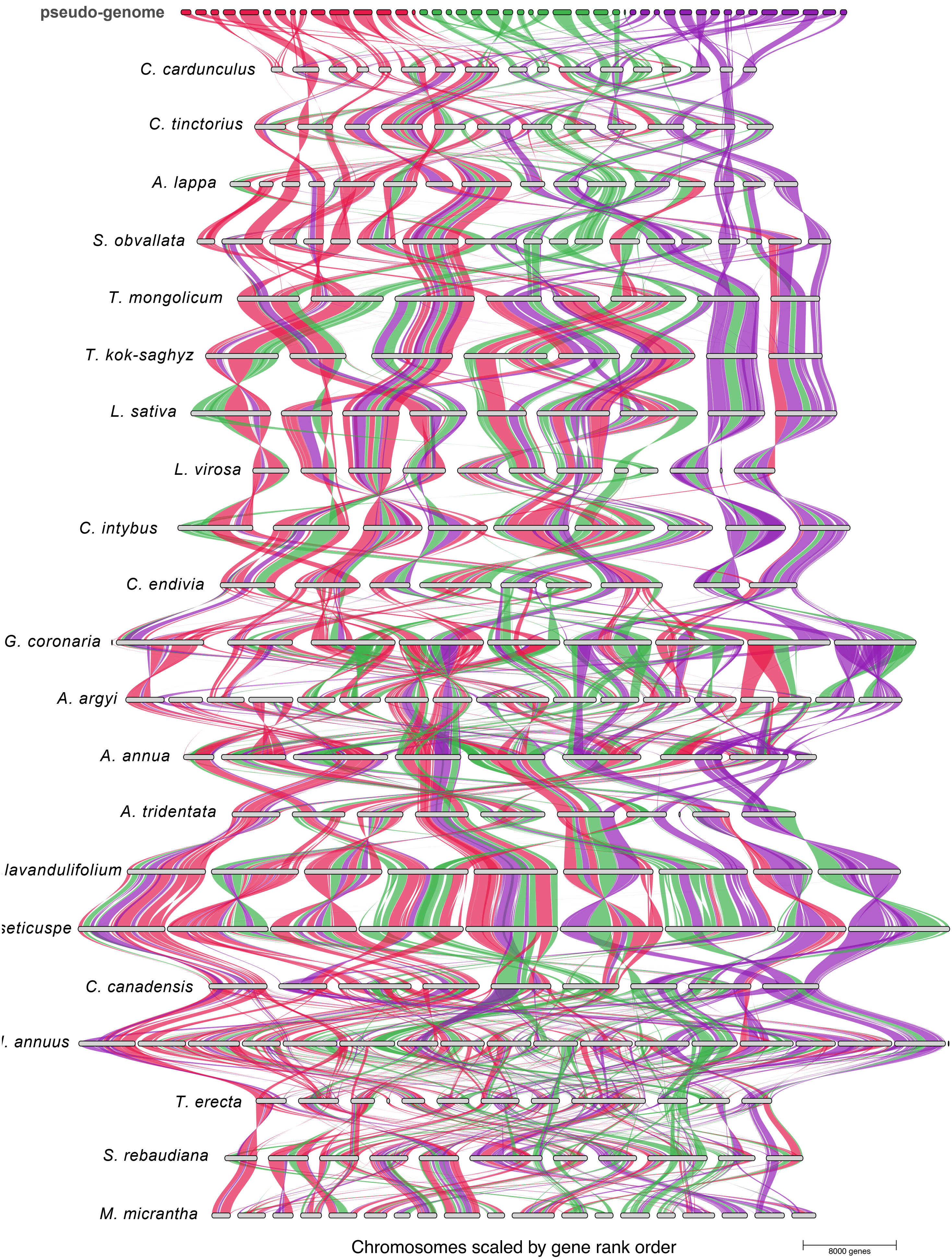

### Fig. S9

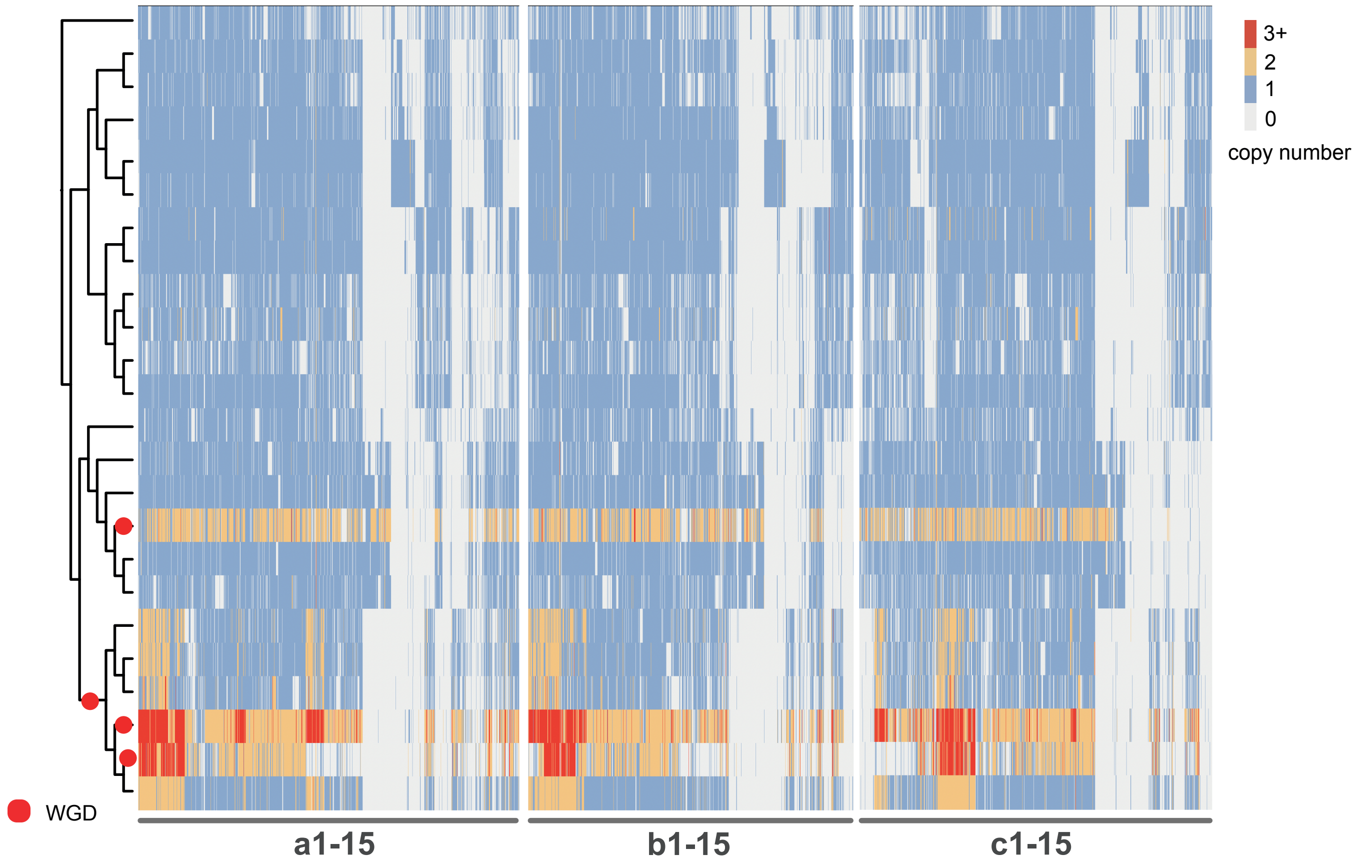

### Fig. S10

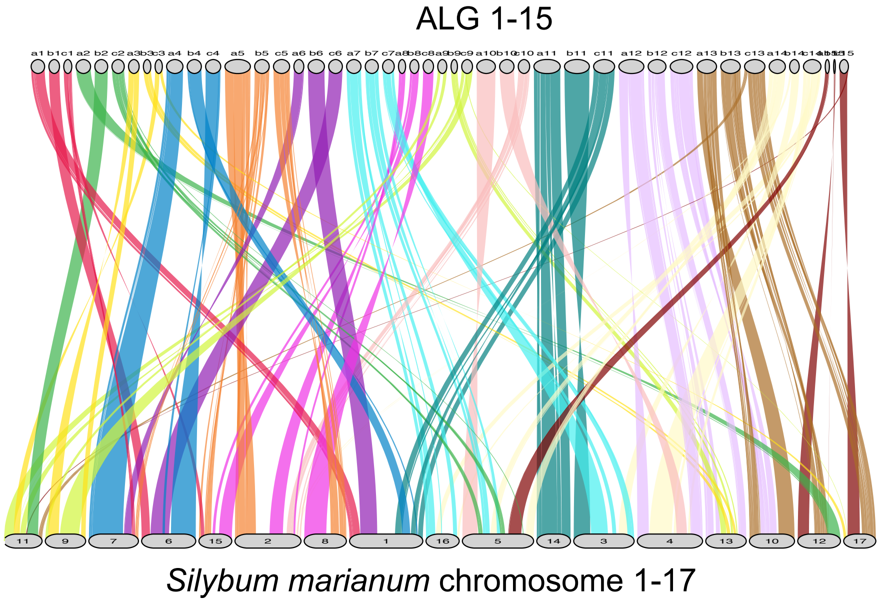

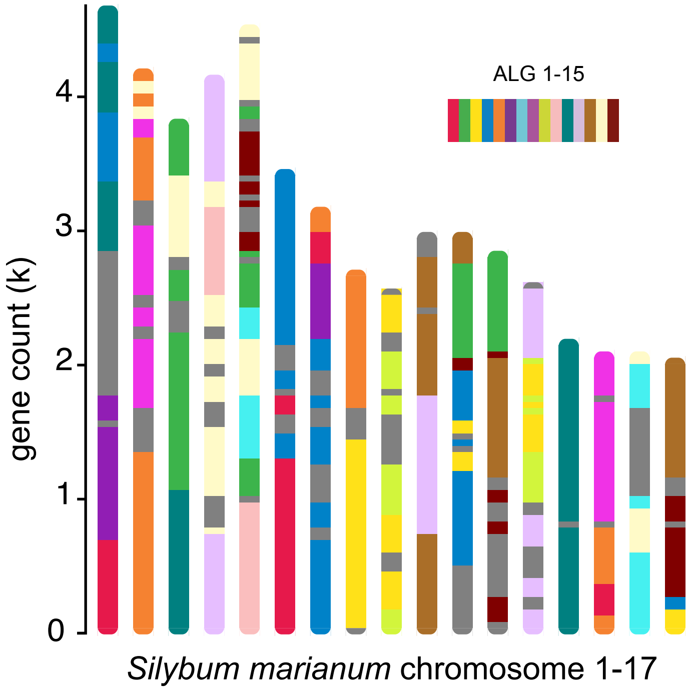

### Fig. S11

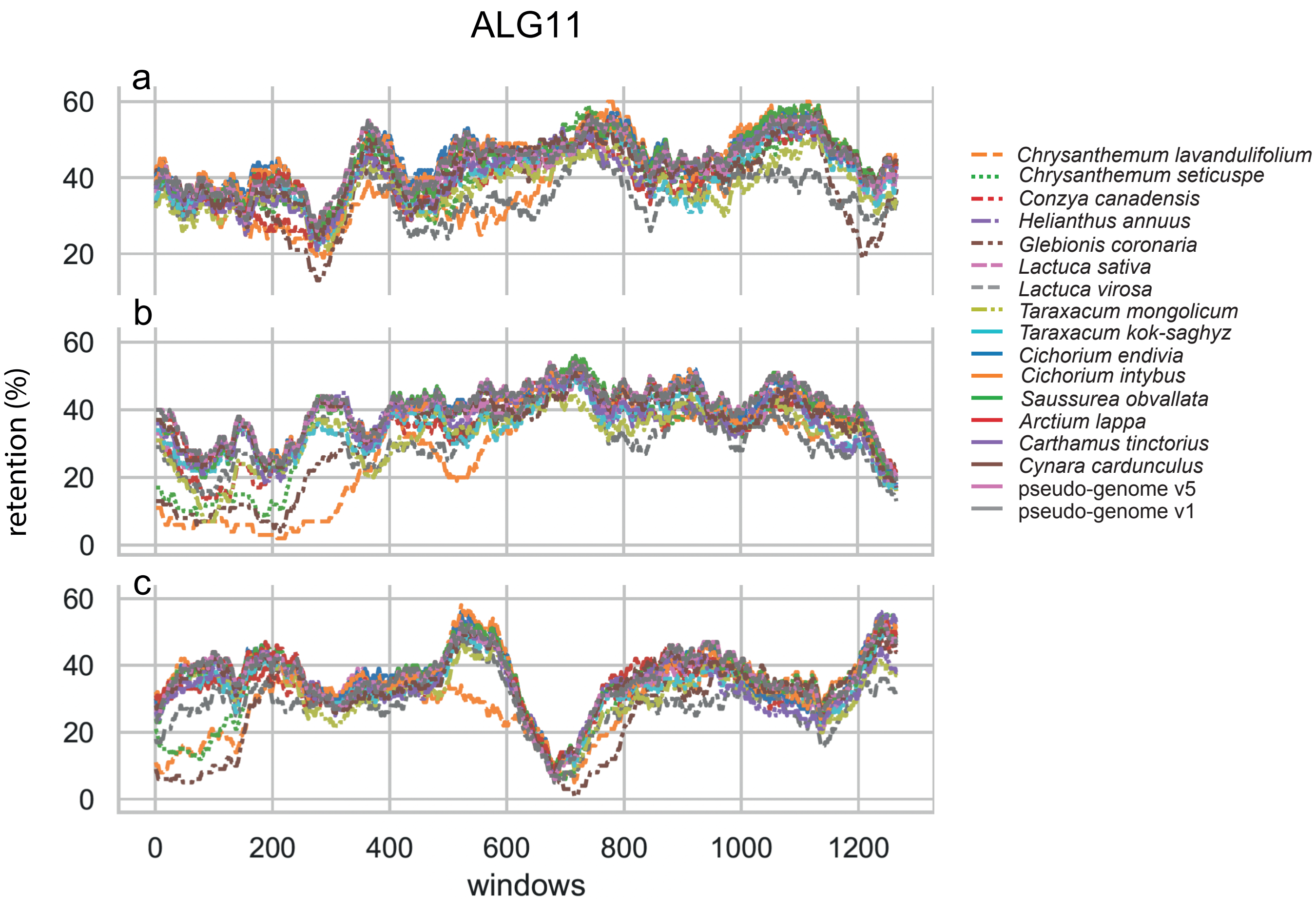

### Fig. S12

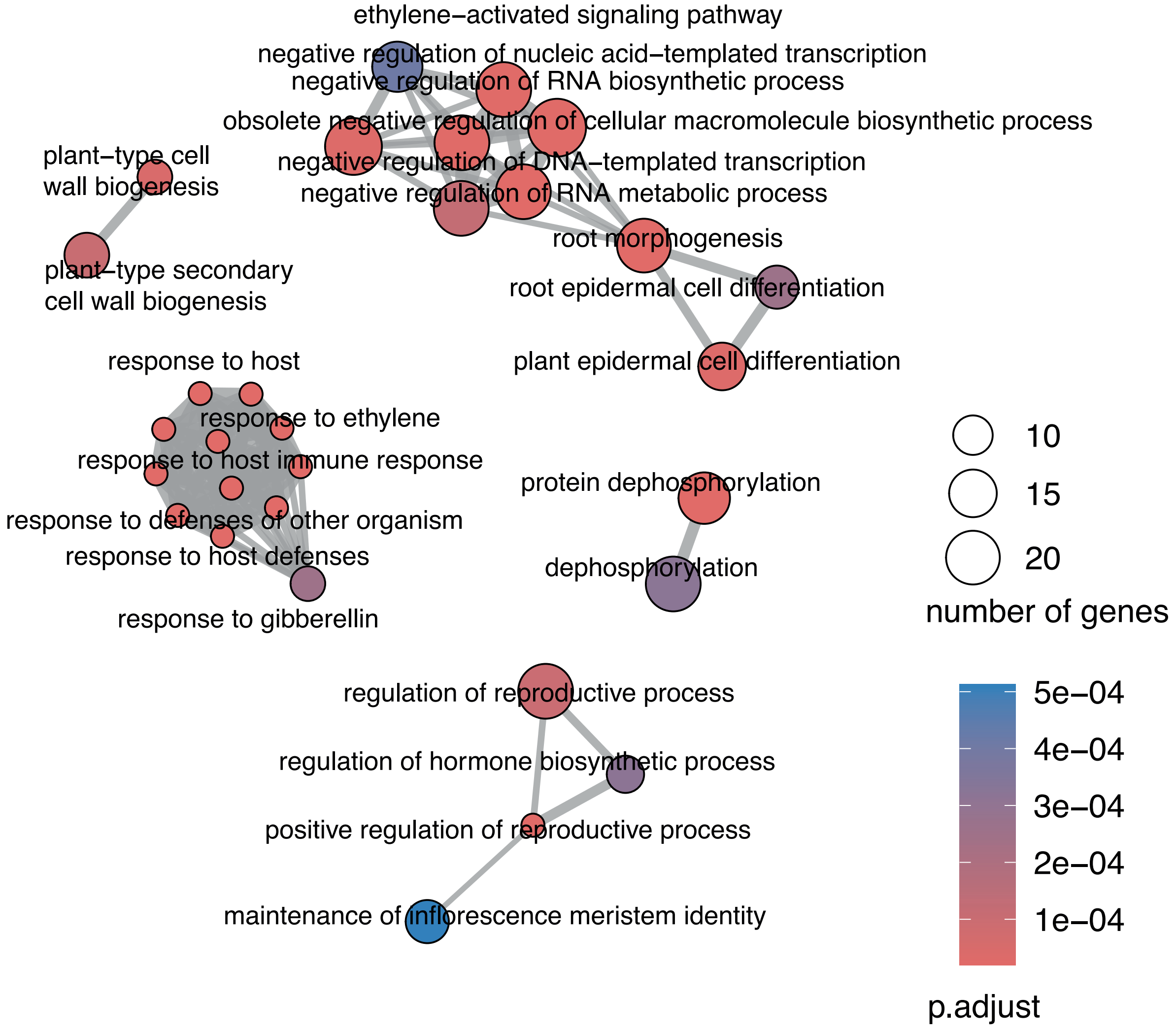
